# *Akkermansia muciniphila* prevents cadmium-induced cognitive impairment through gut microbiome–mediated mechanisms

**DOI:** 10.64898/2026.07.31.740339

**Authors:** Hao Wang, Joe Jongpyo Lim, Jinhua Chi, Haiwei Gu, Julia Yue Cui

## Abstract

*Akkermansia muciniphila* has emerged as a promising next-generation probiotic with beneficial effects on learning and memory, but whether and how it can protect against environmental toxicant-induced cognitive impairment remains unknown. Cadmium (Cd) is a widespread environmental neurotoxicant that disrupts the gut-brain axis and impairs hippocampus-dependent learning and memory, yet effective preventive interventions are lacking. In this study, we discovered that oral supplementation with human fecal microbiome-derived *A. muciniphila* prevented Cd-induced cognitive impairment in mice throughout 9 weeks of oral Cd exposure at a human body burden-relevant concentration. Notably, brain Cd concentrations were not affected by *A. muciniphila* supplementation, indicating that cognitive protection was mediated through gut-brain signaling rather than affecting metal accumulation in the brain. Multi-omics characterization identified coordinated gut-brain pathways underlying this protective effect. *A. muciniphila* preserved Cd-suppressed *Lactobacillus* taxa (*L. crispatus, L. intestinalis, L. taiwanensis*), which positively correlated with cognitive performance, and restored intestinal tight-junction integrity across multiple intestinal sections, particularly the ileum. *A. muciniphila* also normalized Cd-induced cytokine dysregulation in the serum. In addition, colonic branched-chain fatty acids (BCFAs) emerged as candidate gut-brain mediators, with 2-methylpentanoic acid showing a robust negative correlation with cognitive performance. These *A. muciniphila*-mediated changes across gut microbiome, intestinal barrier, systemic cytokines, and microbial metabolites coincided with reversal of Cd-induced hippocampal transcriptional alterations regulating synaptic and vascular signaling. Together, this study identified *A. muciniphila* as a preventive microbiome-based strategy against environmental Cd neurotoxicity in mice, demonstrated that the gut microbial homeostasis can confer cognitive resilience independently of brain toxicant burden, and revealed distinct BCFAs as potential gut-brain mediators of heavy-metal-induced cognitive decline.

## Introduction

Cadmium (Cd) is a naturally occurring heavy metal that has also been extensively used in industrial and commercial applications. Over time, its extensive use and improper disposal have resulted in pervasive environmental Cd contamination. The World Health Organization (WHO) has classified Cd as one of the top ten priority environmental contaminants of major public health concern ^1^. In the general population, Cd exposure occurs primarily through food consumption and cigarette smoking ^2^. Due to its long biological half-life in humans, Cd progressively accumulates in multiple organs, and chronic exposure can result in toxicity affecting liver, kidney, lung, and other systems ^2^.

Cd is considered as an emerging neurotoxicant ^3^. Epidemiological studies have linked Cd exposure to various neurological disorders, including general cognitive decline, Alzheimer’s disease (AD), and Parkinson’s disease (PD) ^4,5^. Consistent with these findings, our previous work demonstrated that Cd exposure at human-relevant levels impairs hippocampus-dependent learning and memory in mouse models ^6–9^. However, the molecular and cellular mechanisms underlying Cd neurotoxicity remain incompletely understood.

Recent studies suggest that the gut microbiome plays a critical role in regulating learning and memory. Prior studies have reported significant gut dysbiosis in AD patients and transgenic AD mouse models ^10–12^. Mechanistically, the gut microbiome can influence cognition through the gut-brain axis, a bidirectional signaling network between the gastrointestinal (GI) tract and the central nervous system (CNS) ^13^. Through these mechanisms, the gut microbiome can modulate cognitive function by regulating neuroactive microbial metabolites, intestinal barrier integrity, and systematic inflammation. For example, short-chain fatty acid, such as butyrate have been shown to improve cognitive function in mice ^10,13^. In contrast, alterations in gut microbiome-derived metabolites and/or pathogenic bacterial products can compromise the intestinal barrier integrity and promote the translocation of inflammatory or neuroactive factors that adversely affect normal CNS function ^14,15^. Building on these recent literature findings on the gut-brain axis, modulation of the gut microbiome has emerged as a potential intervention strategy for improving cognitive impairment. Interventions such as fecal microbiota transplantation (FMT) have shown preliminary benefits in improving learning and memory in AD patients ^16,17^. However, the specific gut taxa responsible for neuroprotective benefits remains unclear.

Notably, the gut microbiome is one of the major targets of Cd toxicity. Epidemiologic studies have found that Cd exposure is associated with significant alterations of fecal microbiome diversity and composition in humans ^18^. Similarly, animal studies demonstrated that Cd exposure can alter the gut microbiome composition, related microbial metabolites, and intestinal integrity ^19,20^. Our recent work was the first to show that Cd-induced gut dysbiosis is associated with Cd-induced hippocampus-dependent learning and memory deficits in mice ^21^. Together, these findings suggest that the gut microbiome contribute significantly to Cd-induced learning and memory impairments. However, whether and how the gut microbiome can be targeted to prevent Cd-induced cognitive impairment remains to be established.

Emerging literature evidence suggests that probiotic, defined as live microorganisms that confer advantageous effects to the host when consumed in adequate amounts ^22^, represent a novel and more targeted approach to improve learning and memory in both animals ^23^ and humans ^24–26^*. Akkermansia muciniphila (A. muciniphila)*, the only currently culturable representative of the phylum *Verrucomicrobia*, is a mucin-degrading commensal bacterium residing predominantly within the mucus layer of the human intestine that has emerged as a promising next-generation probiotics ^27^. It constitutes approximately 3-5% of the gut microbial community ^28,29^ and 1-4% of the fecal microbiota in healthy individuals ^30,31^. In monoculture, *A. municiphila* produces succinate and acetate ^29^, which are cross-fed to butyrogenic bacteria in the intestine, ultimately producing butyrate and other SCFA species ^32,33^. *A. muciniphila* plays a key role in maintaining intestinal barrier integrity by regulating mucus layer thickness, epithelial tight junctions, and host immune responses. For example, altered abundance of *A. muciniphila* has been associated with various intestinal disorders, including ulcerative colitis, inflammatory bowel disease, and acute appendicitis ^34^. *A. muciniphila* has also been shown to promote glucagon like peptide 1 (GLP-1) secretion to improve glucose homeostasis and metabolic disease outcomes ^35^.

Beyond its established role in gut homeostasis, *A. muciniphila* has recently been implicated in the regulation of cognitive function through the gut-brain axis signaling. Recent studies have demonstrated that supplementation with *A. muciniphila* can ameliorate cognitive deficits and attenuate neuropathological features in animal models of neurodegenerative diseases ^36–39^. *A. muciniphila* has also been shown to mechanistically contribute to ketogenic diet-induced seizure protection ^40^. More recently, *A. muciniphila*-derived extracellular vesicles were reported to alleviate colitis-related cognitive deficits by restoring intestinal and blood-brain barrier integrity, attenuating brain inflammation, and remodeling the gut microbiota to preserve synaptic function along the gut-brain axis ^41^.

Given that Cd exposure disrupts gut barrier integrity, promotes systemic inflammatory signaling, and induces gut dysbiosis – all processes in which *A. muciniphila* plays a protective role – we hypothesized that *A. muciniphila* supplementation can protect against Cd-impaired learning and memory through modulating the gut-brain axis. To test this hypothesis, we conducted longitudinal behavioral assessments integrated with multi-omics characterization of the gut microbiome, metabolome, and hippocampal transcriptome, together with intestinal barrier and serum cytokine analyses, in adult male C57BL/6J mice. This study provides the first evidence that microbiome-targeted intervention can confer functional protection against environmental neurotoxicant-induced cognitive impairment through the gut-brain axis.

## 2. Materials and Methods

### Chemicals

Cadmium chloride (CdCl_2_; catalog number: 202980) was purchased from MilliporeSigma (Burlington, MA). Acetic acid was bought from Thermo Fisher Scientific (Fair Lawn, NJ). Propionic acid, isobutyric acid, butyric acid, 2-methylbutyric acid, isovaleric acid, valeric acid, 2-methylpentanoic acid, 3-methylpentanoic acid, isocaproic acid, caproic acid, 2-methylhexanoic acid, 4-methylhexanoic acid, heptanoic acid, hexanoic acid-6,6,6-d3 internal standard, N-tert-Butyldimethylsilyl-N-methyltrifluoroacetamide (MTBSTFA), and methoxyamine hydrochloride were purchased from Sigma-Aldrich (St. Louis, MO). Unless otherwise noted, all other chemicals were purchased from Sigma-Aldrich.

### Animals and exposures

Six-week-old male C57BL/6J mice were purchased from Jackson Laboratories (Bar Harbor, ME) and co-housed (maximum 5 animals per cage) under standard conditions (12-hour light/dark cycle) with ad libitum access to feed (Picolab Rodent Diet 20, LabDiet, St. Louis, MO) and water (tap water purified by reverse osmosis, acidified with 2.4-2.8% HCl, and autoclaved). Beginning at sevenweeks of age, mice received *Akkermansia muciniphila* (BAA-835, obtained from ATCC, Manassas, VA) or vehicle control through oral gavage (10 ml/kg, 1X10^9^ colony-forming units in 0.5 g/L L-cysteine to maintain anaerobic bacterial viability) once daily, five consecutive days per week. After one week of *A. muciniphila* administration, mice were provided either normal drinking water or drinking water containing 3 mg/L Cd (in the form of CdCl_2_) beginning at 8 weeks of age and continuing for 9 weeks **(Fig.1)**. *A. muciniphila* or vehicle treatment was continued throughout the Cd exposure period, for a total treatment duration of 10 weeks. Behavioral tests were conducted before and during the entire exposure period to evaluate the effects of Cd exposure and its interaction with *A. municiphila* on learning and memory. Fresh fecal pellets were collected before *A. muciniphila* treatment (week 0) and at weeks 1, 3, 5, and 9 during the Cd exposure period. In summary, four exposure groups were included in this study: vehicle control (Ctrl), *A. muciniphila* only (AKK), Cd only (Cd), as well as *A. muciniphila* plus Cd co-exposure (AKK+Cd). Mice were euthanized and tissue collection was performed at the end of the study (week 10). The preparation, use, and disposal of hazardous reagents were conducted according to the guidelines set forth by the Environmental Health and Safety Office at the University of Washington. All animal care and experimental procedures were approved by the Institutional Animal Care and Use Committee (IACUC) of the University of Washington and the University of Michigan.

**Figure 1.**
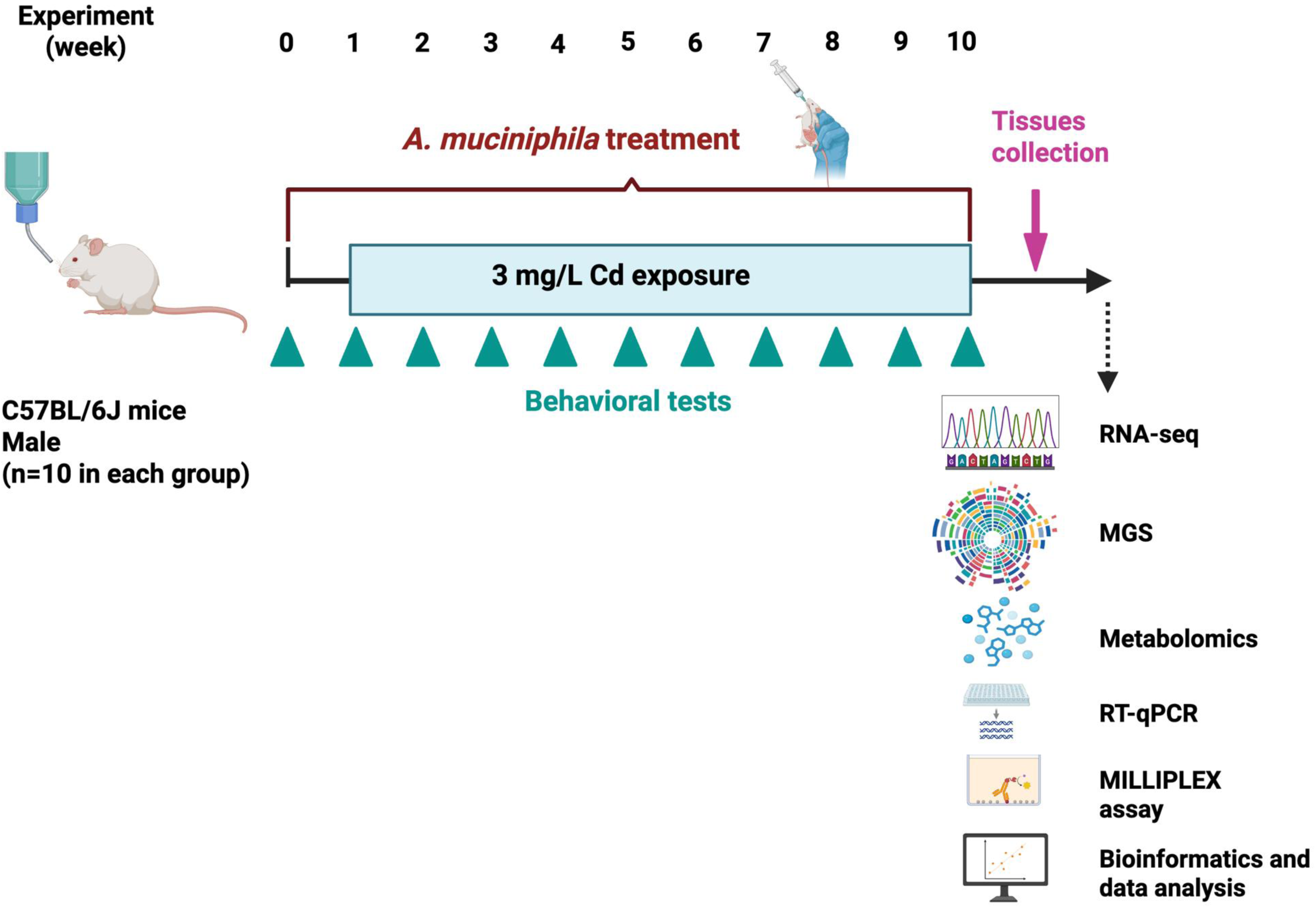
Experimental design of the study. Male C57BL/6J mice were randomly assigned to four groups: Control, *A. muciniphila* (AKK), Cd, and AKK+Cd. *A. muciniphila* was administered by oral gavage beginning 1 week before Cd exposure and continued throughout the experiment. Cd exposure was initiated at week 1 and maintained for 9 weeks through drinking water at 3 mg/L. Behavioral tests were performed longitudinally during the exposure period. At the end of the experiment, tissues were collected for downstream analyses.

### Brain Cd concentration analysis

Brain samples were colleted at the end of the study. The measurement of Cd lelves in the brain were performed by the Environmental Health Laboratory at the University of Washington using inductively coupled plasma mass spectrometry (ICP-MS). All measurement were conducted by using an Agilent 7900 (Agilent Technologies, Santa Clara, California) which has a detection limit of 0.08ng/ml.

### Open field test

The open field test was used to assess locomotor activity and anxiety. The test was performed before and at the end of exposure following our previous protocol ^9,21^. In general, mice were placed into a 10 inches (width) × 10 inches (depth) × 16 inches (height) TruScan Photo Beam Tracking arena (Coulbourn Instruments, Whitehall, PA) with clear Plexiglas sidewalls, and their movement was monitored with two sets of infrared breams spaced 0.6 inches apart during the test. Animals were allowed to freely explore the arena without pre-habituation for 20 min, and data were collected by TruScan 2.0 software (Coulbourn Instruments). The total move distance, total move time, and average speed were used to assess locomotor activity. In addition, time spent in the margin and center zones, distance traveled in the margin and center zones, and the number of center entries were used to examine anxiety-like behavior.

### Novel object location test

The novel object location test was used to assess the hippocampus-dependent spatial memory following treatments ^9^. The assay was performed every other week as previously described ^9^. Briefly, mice were placed in an open field arena (Coulbourn Instruments) with two identical objects placed in two different corners. During the training session, mice were allowed to freely explore the two objects for 5 min and were then returned to their home cages. To exclude potential preference for a specific location, alternating corners were used for object presentation. The testing session was performed 1 h after the training session, and animals were returned to the arena with the same two objects, with one object remaining in its original location and the other moved to a novel location. The time animals spent actively investigating each object during the training and testing sessions was recorded by cameras and quantified by an experimenter blinded to the animal’s treatment.

For each test session the discrimination index (DI) was calculated for each mouse as DI = (T_novel_ -T_original_)/(T_novel_ + T_original_) where T_novel_ and T_original_ represent the time spent exploring the object in the novel location and the original location, respectively. DI values range from -1 to 1, with 0 indicating chance-level performance, positive values indicating preferential exploration of the object in the novel location, and negative values indicating preference for the object in the familiar location. DI values were analyzed using a linear mixed-effects model (LMM) with Group (Ctrl, AKK, Cd, and AKK+Cd) and Time (week 3, 5, 7, and 9), and their interaction included as fixed factors. Individual mouse ID was included as a random intercept to account for within-subject correlations across time points. Models were fitted using the *lme* package (v 2.0.1) with restricted maximum likelihood (REML) estimation. Missing values were handled implicitly by the LMM under the missing-at-random assumption. Estimated marginal means and post-hoc pairwise comparisons between groups at each timepoint were obtained using the *emmeans* package (v 2.0.2). P-values from multiple comparisons were adjusted using the Benjamini-Hochberg (BH) false discovery rate (FDR) correction, applied separately within each timepoint. All analyses were performed in R (v 4.5.3).

### Fecal DNA extraction and shallow metagenomic shotgun sequencing

DNA was extracted from 100 - 200 mg fecal samples by using an AllPrep^®^ PowerFecal DNA/RNA Kit (Qiagen, Hilden, Germany) according to the manufacturer’s protocol. DNA concentration was quantified using a Qubit 2.0 Fluorometer (Life Technologies, Grand Island, NY). The integrity and quality of DNA samples were confirmed by using an Agilent 2100 Bioanalyzer (Agilent Technologies Inc., Santa Clara, CA). Shallow metagenomic shotgun sequencing was performed by Diversigen (New Brighton, MN). Sequencing reads were aligned to a curated database containing all representative genomes in RefSeq for bacteria with additional metagenomically assembled genomes (MAGs) and cell-cultured genomes. Only high-quality MAGs (Completeness > 90% and Contamination < 5% via checkm) were considered. Reads were aligned to all reference genomes at 97% identity. Each input sequence was compared to every reference sequence in the Diversigen Venti database using fully gapped alignment with BURST. Ties were resolved by minimizing the total number of unique Operational Taxonomic Units (OTUs). For taxonomy assignment, each input sequence was assigned to the lowest common ancestor that was consistent across at least 80% of all reference sequences tied for best hit. Taxonomic annotations were based on Genome Taxonomy Database (GTDB r95). Samples with fewer than 10,000 sequences were excluded from further analysis. OTUs accounting for less than one-millionth of all strain-level markers and those with less than 0.01% of their unique genome regions covered (and < 0.1% of the whole genome) at the species level were also removed.

To account for the compositional nature of microbiome count data, taxonomic abundance profiles were transformed using centered log-ratio (CLR) transformation prior to downstream statistical analyses **(Supplementary Table 2)**. CLR-transformed abundance values were used for longitudinal modeling and visualization. Heatmaps and boxplots were plotted using ComplexHeatmap (V.2.25.2)^42^ and *ggplot2* (version 4.0.0)^43^.

### Longitudinal analysis of microbial taxa and L2/L3 KEGG enzyme functional features

Longitudinal differential abundance analysis of the gut microbiome was performed using linear mixed effects modeling to account for repeated measurements within individual mice. For the initial screening, each microbial taxon was analyzed using the model **Abundance** ∼ **Group × Time + (1**∣**Subject_ID)**. In this model, **Group** included four types of treatments [Control, AKK (*A. muciniphila)*, Cd, and AKK+Cd], **Time** was treated as a categorical variable representing the sampling timepoints, and **Subject_ID** was included as a random intercept to account for within-mouse correlation across repeated observations. This framework was used to identify taxa exhibiting overall group differences and/or treatment-dependent temporal abundance patterns.

For each taxon, the global Group x Time interaction term was extracted from the linear mixed effects model to determine whether longitudinal trajectories differed across the four different groups. Taxa with a nominally significant global interaction effect (p < 0.05) were retained as candidate taxa for downstream analyses. Because this step was intended to identify taxa showing overall treatment-related differences in longitudinal trajectories, selection was based solely on the global Group × Time interaction *p* value rather than on individual timepoint-specific contrasts. This approach allowed us to capture taxa with consistent or distributed temporal effects across the exposure period, while reducing reliance on isolated week-specific fluctuations. Using this criterion, 65 taxa were identified for subsequent follow-up analyses.

For these 65 candidate taxa, targeted post hoc contrasts were then performed to assess differences between the Cd and AKK+Cd groups during the active Cd exposure phase (weeks 1, 3, 5, and 9). Estimated marginal means (EMMs) were calculated for each taxon, and pairwise comparisons were restricted to these biological relevant exposure time points. Baseline (before *A. muciniphila* treatment) and week 0 (one week after *A. muciniphila* treatment) samples were retained in the mixed-effects models to improve estimation of subject-specific intercepts and overall longitudinal structure but were not included in the exposure-phase *post hoc* testing because Cd exposure had not yet begun. To further characterize differences between the Cd and AKK+Cd groups during the active Cd exposure phase, both the overall Group × Time interaction and week-specific Cd vs. AKK+Cd contrasts were evaluated for each candidate taxon. Benjamini-Hochberg FDR correction was applied to both the overall interaction tests and the week-specific post hoc comparisons. Only taxa with FDR-adjusted p < 0.05 were considered statistically supported in the final interpretation **(Supplementary Table 3)**.

For the *Lactobacillus* taxa analysis, the Cd and AKK+Cd groups were compared across six time points (baseline and week 0-9). To test whether Cd and AKK+Cd groups exhibited divergent abundance trajectories, we fitted **Abundance** ∼ **Baseline_Abundance + Group x Time + (1**∣**Subject_ID)**, where **Time** was a categorical factor (weeks 1, 3, 5, and 9 after Cd exposure), and Subject_ID was a random intercept to account for repeated measurements. The group x time interaction term was used to test whether temporal trajectories differed between groups. To assess the direction and magnitude of abundance changes within each group, change from week 0 was calculated as ΔCLR (CLR abundance at time t − CLR abundance at week 0). Separate models [Δ**CLR ∼ Time + (1 | Subject_ID)**] were fitted for each group, with Time treated as a categorical variable (weeks 1, 3, 5, and 9). Estimated marginal means at each time point were tested against zero using t-tests. As supplementary evidence, between group differences in ΔCLR at each time point were assessing using ΔCLR ∼ Group x Time + (1 | Subject_ID). Pairwise comparison between Cd and AKK+Cd groups were extracted at each time point using the emmeans package. Linear mixed-effects models were fitted using the lme4 (v1.1-35.1) ^44^ and lmerTest (v3.1-3) ^45^ packages in R (v4.3.0), with restricted maximum likelihood (REML) estimation. *P*-values were calculated using Satterthwaite’s method for degrees of freedom approximation. Estimated marginal means and pairwise contrasts were calculated using the *emmeans* package (v1.8.9) ^46^. All p-values were adjusted for multiple comparisons using the Benjamini–Hochberg FDR adjustment procedure. Statistical significance was defined as FDR-adjusted p < 0.05. CLR-transformed reads were used to further investigate group-wise change in abundance patterns for each time point and plotted using ggplot2 (v 4.0.1) _43,_

Predicted functional profiles annotated to the KEGG enzyme (EC) hierarchy were generated by Diversigen (New Brighton, MN). For the functional analysis, we used the same methods as for the longitudinal taxa analysis, applying them to CLR-transformed counts of predicted KEGG enzyme features at level 2 and level 3 **(Supplementary Table 4)**.

### Correlation analysis between gut microbiota and behavioral performance

Repeated-measures correlation analysis between CLR abundance and the discrimination index (DI) from the novel object location test was performed to investigate longitudinal association between candidate taxa and spatial memory. Samples from weeks 3, 5, and 9 after Cd exposure in the Cd and AKK+Cd groups were included in this analysis. The analysis was performed using the rmcorr package (v0.5.6) in R, with Subject_ID specified separately. To ensure statistical reliability, each taxon was included in the analysis only if it met the following criteria after removal of missing values: (1) non-zero variance in CLR abundance, (2) at least 2 subjects with complete data, (3) at least 4 total observations across all subjects and time points, and (4) at least two time points contributed by each included subject. These criteria were applied to ensure that sufficient repeated observations were available to estimate within-subject correlations. For each qualifying taxon, the repeated-measures correlation coefficient (r), degrees of freedom, and nominal p value were extracted. Multiple testing correction across all tested taxa was performed using the Benjamini-Hochberg FDR adjustment.

### Metagenomic alignment to Akkermansia reference genomes

Murine (strain:139 ^47^, Assembly ASM431956v1) and human (ATCC BAA-835, Assembly ASM1750414v1) isolates of *Akkermansia muciniphila* were retrieved from RefSeq. Stool metagenomes in all treatment groups were aligned to both *Akkermansia* strains to directly compare the abundance of murine and human Akk abundance in the gut. The reference index was created from the *Akk* isolates using bowtie2-build. Metagenomic fastq files were directly aligned to the reference index using bowtie2 (v 2.5.5) ^48^. BAM files were then created using samtools (v 1.21) ^49^. The successfully mapped reads were then aggregated by genome. For each sample, the total mapped reads for each genome were normalized to the total reads. The log ratio between BAA-835 and strain:139 was taken as a measure of human to mouse *A. muciniphila* relative abundance (i.e., positive value indicates higher BAA-835 over murine *A. muciniphila*).

### RNA isolation

Total RNA was isolated from hippocampal and intestinal tissues colleted from mice in all four groups after exposure using the RNA-Bee reagent (Tel-Test Inc, Friendswood, TX). RNA concentrations were quantified by using a NanoDrop 1000 Spectrophotometer (Thermo Scientific, Waltham, MA) at 260 nm. The quality of RNA was evaluated by formaldehyde-agarose gel electrophoresis by visualizing the 28S and 18S rRNA bands under UV light and Agilent 2100 Bioanalyzer by Novogene (Sacramento, CA). RNA sequencing was performed by Novogene using 4 biological replicates.

### RNA-Seq of hippocampal transcriptome

FASTQ files with paired-end sequencing reads were mapped to the mouse genome (UCSC mm10) using HISAT2 (Hierarchical Indexing for Spliced Alignment of Transcripts) ^50^. The resulting SAM (sequence alignment/map) files were converted to their binary form and sorted using SAMtools (v 1.2) ^49^. Transcript abundances were estimated with featureCounts (part of the Subread package, v 1.5.3) ^51^ using the Gencode mouse version 25 (vM25) gene transfer format (GTF). Differential expression analysis was performed using Cuffdiff ^52^, with FDR-adjusted p < 0.05 considered as statistically significant **(Supplementary Table 5)**. Gene ontology enrichment analysis was conducted using the topGO package (v 2.52.0) ^53^ in R. Count data were normalized to transcript per million (TPM). Principal component analysis (PCA) was performed on the filtered TPM normalized counts (above 1^st^ quantile of standard deviation) using the princomp function in R (v 4.5.3).

### RT-qPCR quantification

Total RNA isolated from intestinal tissues and intestinal contents was reverse transcribed into cDNA using a high-capacity cDNA Reverse Transcription Kit (Life Technologies, CA). The resulting cDNA products were then amplified by qPCR, using the Sso Advanced Universal SYBR Green Supermix in a Bio-Rad CFX384 Real-Time PCR detection system (Bio-Rad, Hercules, CA). *A. muciniphila* abundance was quantified by qPCR using primers (forward: 5′-TAGTCCAGGGCATGCAGAAA-3′; reverse: 5′-ATCCCTGAAAACAACCGTGC-3′) originally designed against the *A. muciniphila* ATCC BAA-835 reference genome. Primer sequences for tight-junction related genes and housekeeping genes are provided in the **Supplementary Table 1**. Gene expression data were normalized to the housekeeping gene using the ΔΔCq method and were expressed as % of the housekeeping gene **(Supplementary Table 6)**.

### Serum cytokine levels quantification

Serum cytokine levels were quantified using the MILLIPLEX^®^ Mouse High Sensitivity T Cell Magnetic Bead Panel assay (Millipore Sigma, Burlington, MA. Catalog number: MHSTCMAG-70KPMX) following the manufacturer’s instructions. Briefly, quality controls, standards, and serum samples were prepared in triplicate and added into a 96-well plate pre-coated with magnetic beads conjugated with capture antibodies specific to target cytokines. After overnight incubation at 4 °C, the plates were washed by using a wash buffer to remove unbound components. Detection antibodies were added into each well and incubated for one hour at room temperature, followed by the incubation with streptavidin-phycoerythrin (PE) for 30 min. After 3 times of gentle washes, the beads were resuspended in assay buffer and analyzed using the Belysa^®^ Immunoassay Curve Fitting Software.

### Short-chain fatty acids analysis

50 mg of each tissue sample was homogenized with 20 μL hexanoic acid-6,6,6-d_3_ (internal standard; 200 µM in H_2_O), 20 μL sodium hydroxide solution (NaOH, 0.5 M in water), and 480 μL methanol (MeOH). Afterward, 400 μL MeOH was added, and the pH of the mixture was approximately 10. Upon storage at -20°C for 20 min and centrifugation at 21,694 g for 10 min, 800 μL of supernatant were collected. Samples were then evaporated to dryness, reconstituted in 40 μL of methoxyamine hydrochloride in pyridine (20 mg/mL), and stored at 60°C for 90 min. Afterward, 60 μL of N-Methyl-N-tert-butyldimethylsilyltrifluoroacetamide was added and stored at 60°C for 30 min. Each sample was then vortexed for 30 s and centrifuged at 21,694 g for 10 min. Finally, 70 μL of supernatant was collected from each sample for GC-MS analysis. For serum/intestinal content samples, 20 μL of each sample was mixed with 30 μL aqueous NaOH (0.1M in water), 20 μL IS (hexanoic acid-6,6,6-d_3_; 200 µM) and 430 μL MeOH in a 1.5 mL Eppendorf tube. The pH value for the mixture was 9. Samples were then vortexed for 10 s and stored under -20 °C for 20 min. After centrifugation at 14,000 RPM for 10 min at 4 °C, 450 μL supernatant was removed into a new Eppendorf tube. The samples were dried under vacuum at 37 °C for 120 min using a CentriVap Concentrator (Labconco, Fort Scott, KS). Each sample was first derivatized with 40 µL of methoxyamine hydrochloride solution in pyridine (MeOX, 20 mg/mL) under 60 ℃ for 90 min. Next, 60 µL of MTBSTFA was added, and the mixture was incubated under 60 ℃ for 30 min. Then the sample was vortexed for 30 s, followed by centrifugation at 14,000 rpm for 10 min. Finally, 70 µL supernatant was collected into a new glass vial for GC-MS analysis. GC-MS experiments were performed using an Agilent 7820A gas chromatography system coupled to an Agilent 5977B mass spectrometer (Agilent Technologies, Santa Clara, CA). Chemical derivatives in the samples were separated using an HP-5 ms capillary column coated with 5% phenyl-95% methylpolysiloxane (30 m×250 µm i.d., 0.25 µm film thickness, Agilent Technologies). 1 µL of each sample was injected, and the solvent delay time was set to 5 min. The initial oven temperature was held at 60 ℃ for 1 min, ramped up to 325 ℃ at a rate of 10 ℃/min, and finally held at 325 ℃ for 10 min. Helium was used as the carrier gas at a constant flow rate of 20 mL/min through the column. The temperatures of the front inlet, transfer line, and electron impact (EI) ion source were set at 250 ℃, 290 ℃, and 230 ℃, respectively. The electron energy was -70 eV, and the mass spectral data were collected in the full scan mode (*m/z* 30-600). Agilent MassHunter Workstation Software Quantitative Analysis (B.09.00) was used to process the GC-MS data for compound identification, peak picking, and quantification. The signal-to-noise ratio (S/N) was set to S/N=3. The retention time and quantification mass for each SCFA were determined using its chemical standard. The concentrations of SCFAs in biological samples were calculated using the calibration curves constructed from the corresponding SCFAs standards **(Supplementary Table 7)**.

Each metabolite was analyzed independently using two-way ANOVA, with Cd exposure and *A. muciniphila* treatment as the main factors. For fold-change visualization, log_2_ ratios relative to the control group were calculated for each SCFA. Spearman’s rank correlation analysis was performed to evaluate associations between discrimination index and the levels of selected metabolites, with multiple-testing correction using the Benjamini-Hochberg FDR adjustment method. All statistical analyses and visualization were performed in R (v 4.5.3) using base statistical analysis functions and the *ggplot2* package.

### Statistical analysis

Statistical analyses were performed using GraphPad Prism software (GraphPad Software Inc, La Jolla, CA) or R (v 4.5.3) with the indicated packages. Data are presented as mean ± SEM unless otherwise stated. Microbiome taxonomic and functional analyses were performed on CLR-transformed abundance data using mixed-effects modeling, with false discovery rate correction applied for multiple comparisons where appropriate. RNA-seq differential expression analysis was performed using DESeq2 (v 1.48.2), with Benjamini-Hochberg-adjusted p < 0.05 considered statistically significant. SCFA concentrations and cytokine levels were analyzed using two-way ANOVA followed by post hoc multiple-comparison testing when appropriate. Correlation analyses were performed using repeated-measures correlation or Spearman’s rank correlation, as specified for each analysis. Statistical significance was defined as p < 0.05 or FDR-adjusted p < 0.05, unless otherwise stated. * *p* < 0.05; ** *p* < 0.01; *** *p* < 0.001.

## Results

### *A. muciniphila* supplementation protected mice from Cd-induced hippocampus-dependent learning and memory deficits

To investigate the effects of *A. muciniphila* on Cd-induced hippocampus-dependent learning and memory deficits, C57BL/6J mice were pretreated with *A.muciniphila* via oral gavage (once daily, five consecutive days per week) for one week prior to Cd exposure (3 mg/L, through drinking water). The bacterial treatment was continued throughout the entire Cd exposure period, lasting a total 10 weeks **(Fig. 1)**. Body weight was monitored over the 10-week period, and no significant differences were observed across any exposure group **(Fig. 2A)**. We also quantified the *A. muciniphila* DNA abundance in the large intestinal contents by qPCR and found an increase in its levels in the samples of AKK+Cd groups as compared to the Cd group, as well as a non-significant increasing trend in the AKK group as compared to the controls **(Fig. 2B),** indicating successful establishment of *A. muciniphila* in supplemented mice. To further confirm colonization specificity, we performed strain-resolved metagenomic alignment to distinguish the gavaged human-derived strain (ATCC BAA-835) from the endogenous murine *A. muciniphila*. Fecal metagenomes were aligned to both ATCC BAA-835 and the murine isolate (Strain 139) **(Fig.S1)**, and the log ratio of reads mapped to each genome was used as a measure of relative strain abundance. Both the AKK and AKK+Cd groups maintained consistently positive ratios following gavage, whereas Control and Cd groups remained negative throughout the study period **(Fig. 2C)**, confirming successful colonization of ATCC BAA-835 in supplemented mice.

**Figure 2.**
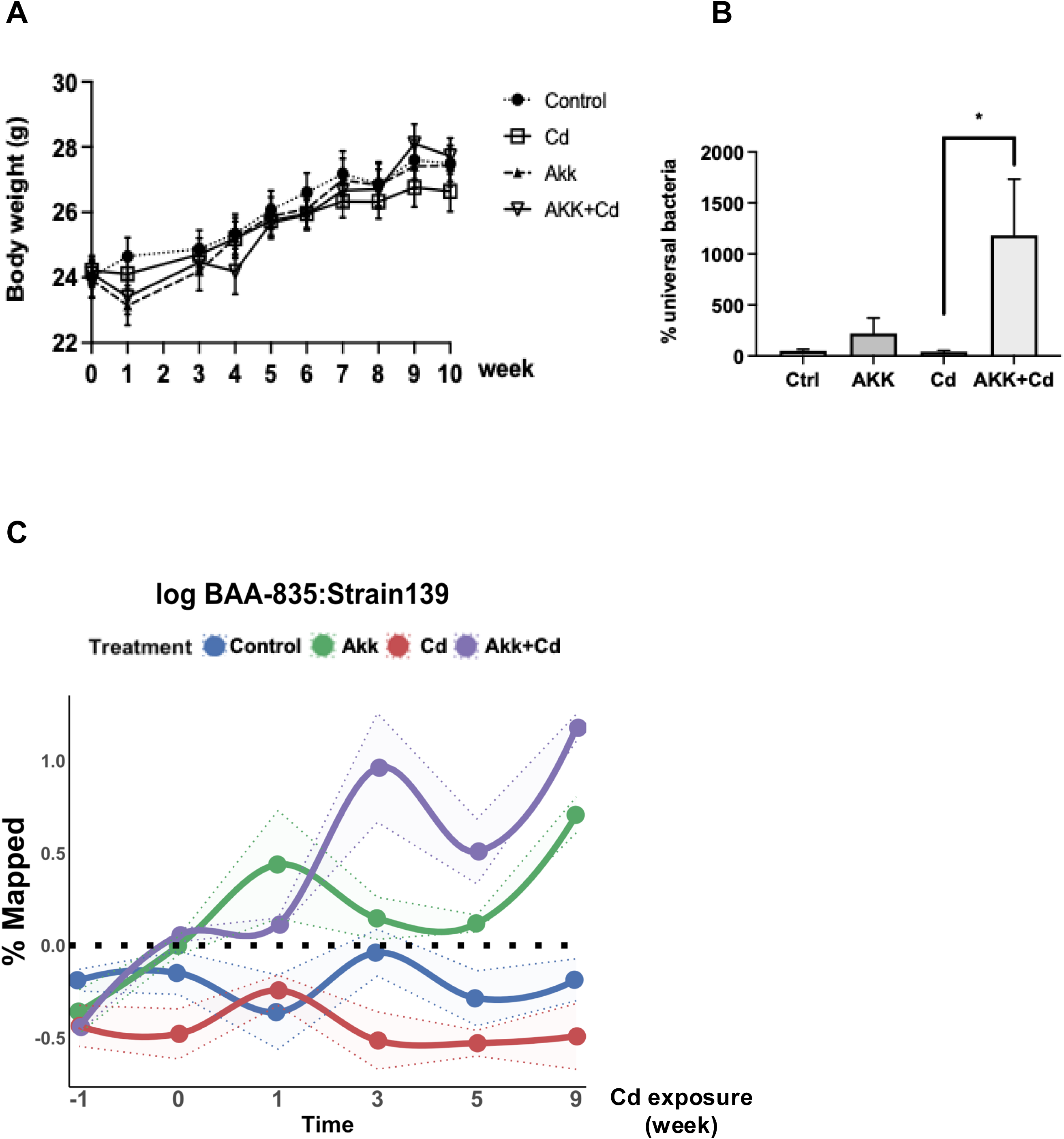
Neither *A. muciniphila* treatment nor Cd exposure affected body weight and increased fecal *A. muciniphila* abundance confirmed successful bacterial colonization. **A.** Weekly body weight measurements in the four groups throughout the whole experimental period. Neither *A. muciniphila* treatment nor Cd exposure induced significant body weight changes. n=9-10 per group. **B.** Relative abundance of *A. muciniphila* in Large intestinal contents was significantly increased in the AKK-treated groups. **C.** Fecal metagenomes from the four-treatment group were aligned to both the human-derived A. muciniphila isolate (ATCC BAA-835) and the murine-derived isolate (Strain 139) using Bowtie2. The log ratio of mapped reads (BAA-835:Strain 139) was calculated for each sample as a measure of relative abundance of the human versus murine *A. muciniphila* strain. Positive values (above the dashed line) indicate predominance of BAA-835, whereas negative values indicate predominance of the endogenous murine strain. Time −1 represents baseline prior to any treatment; Time 0 represents one week after *A. muciniphila* gavage; Time 1, 3, 5, and 9 represent weeks following Cd exposure. n = 4 in each group. Data are presented as mean ± SEM. * *p* < 0.05.

To evaluate the onset of hippocampus-dependent learning and memory deficits, a bi-weekly one-hour Novel Object Location (NOL) test was performed throughout of the whole exposure period **(Fig. 3A)**. During the training sessions, animal across all experimental groups spent similar amount of time exploring each location, indicating no inherent preference for either object or location (data not shown). During the testing session **(Fig. 3B)**, at 3 weeks of Cd exposure, all groups demonstrated intact spatial memory by spending significantly more time exploring the object in the novel location (C) relative to the object in the original location (A). However, by week 5 of Cd exposure, the Cd-only group began to exhibit memory impairment, as evidenced by failure to distinguish between objects in the novel and original locations. At weeks 7 and 9, the Cd group continued to show impaired spatial memory, whereas the AKK+Cd group maintained their ability to discriminate between locations throughout the entire exposure period, indicating that *A. muciniphila* supplementation successfully prevented mice from Cd-induced cognitive deficits.

**Figure 3.**
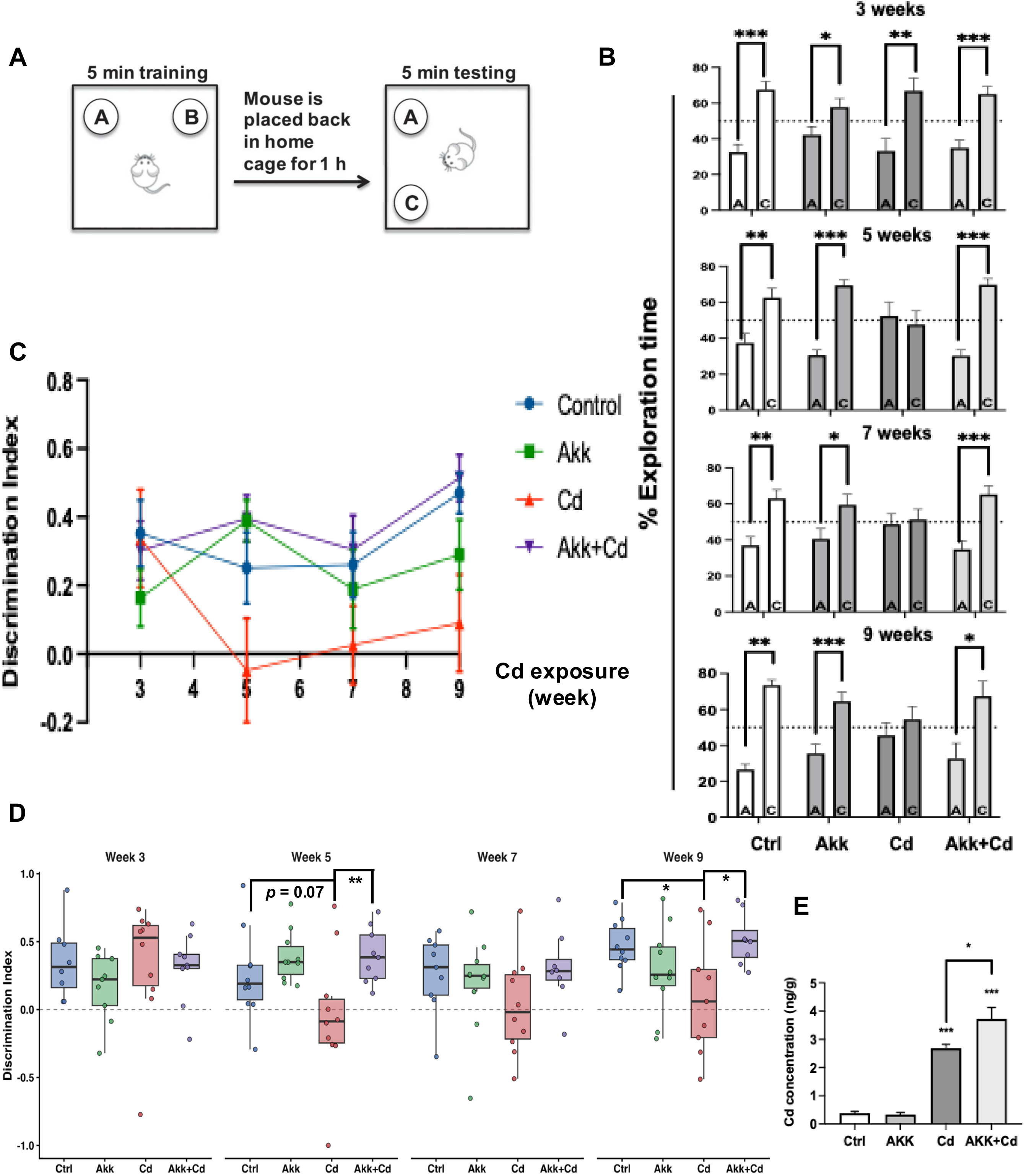
*A. muciniphila* prevented Cd-induced hippocampus-dependent spatial memory impairment in mice. **A.** Schematic illustration of the novel object location (NOL) test. During the 5-min training session, mice were allowed to explore two identical objects placed at locations A and B. After a 1-h interval in the home cage, mice were returned to the arena for a 5-min testing session, in which one object remained at the original location (A) and the other was moved to a novel location (C). **B.** Percentage of exploration time spent investigating the object in the original location or the novel location during the NOL test at weeks 3, 5, 7, and 9. At week 3, all groups showed intact spatial memory by spending significantly more time exploring the object in the novel location. From week 5 onward, mice in the Cd group failed to distinguish between the familiar and novel locations, whereas mice in the AKK+Cd group maintained a clear preference for the object in the novel location. **C.** Longitudinal changes in discrimination index (DI) across the four groups. **D.** Group comparisons of DI at weeks 3, 5, 7, and 9. The Cd group showed reduced DI at later time points, whereas the AKK+Cd group remained comparable to controls. **E.** Brain Cd concentrations measured at the end of the study. n = 8-10 in behavioral test results. n = 4 in Brain Cd measurement. Data are presented as mean ± SEM in **B**, **C**, and **E**; box plots in **D** show the median, interquartile range, and individual data points. * *p* < 0.05; ** *p* < 0.01; *** *p* < 0.001.

We further analyzed the longitudinal changes of Discrimination Index (DI) of the NOL test **(Fig. 3C)** across the four groups using a linear mixed-effects model. The results **(Fig. 3D)** revealed a significant main effect of Group on DI (F = 5.37, *p* = 0.002), while neither the main effect of Time (F = 1.37, *p* = 0.255) nor the Group × Time interaction (F = 1.36, *p* = 0.211) reached statistical significance. At week 3, all four groups showed comparable DI values with no significant differences. Cd exposure induced a progressive decline in cognition. By week 5, Cd-exposed mice showed a substantial reduction in DI, with values falling to near-chance level, while control, AKK, and AKK+Cd groups consistently maintained DI values. At this timepoint, Cd-only mice exhibited significantly lower DI than the AKK+Cd group and showed a strong trend toward reduced DI compared with controls (*q* = 0.077). The trend of Cd-induced deficits remained but did not reach statistical significance at week 7, likely reflecting the increased within-group variability observed at this timepoint. This trend continued by week 9, the deficits were fully re-established, with Cd mice showed significantly lower DI than both the control and AKK+Cd groups. Importantly, AKK+Cd mice were statistically indistinguishable from controls at every timepoint, indicating that *A. muciniphila* supplementation prevented the Cd-induced deficit rather than producing only a partial rescue effect. *A. muciniphila* administration alone had no effect on DI, confirming that the supplementation itself did not alter recognition. Together, these results demonstrate that Cd exposure produces a progressive impairment in hippocampus-dependent spatial working memory that emerges by week 5 and is stably established by week 9, and that *A. muciniphila* supplementation prevented this impairment.

Because cognitive performance can be confounded by alterations in locomotor activity and anxiety, we also performed an open-field test at week 10 (9 weeks into Cd exposure). As shown in **Fig. S2**, neither treatment significantly affected locomotor activity or anxiety-like behavior in mice.

### *A. muciniphila* supplementation preserved cognitive function independently of brain Cd accumulation

Having established that *A. muciniphila* supplementation protects against Cd-induced cognitive deficits, we next explored the mechanisms underlying this protection. We first hypothesized that the protective effect might be induced by reduced Cd accumulation in the brain. To investigate this possibility, we quantified brain Cd concentrations in mice by ICP-MS. Two-way ANOVA revealed significant main effects of Cd exposure and AKK treatment, along with a significant Cd ×AKK interaction (Cd: *p* < 0.0001; AKK: *p* = 0.041; interaction: *p* = 0.027) **(Fig. 3E)**. Post hoc Tukey testing confirmed that both the Cd and AKK+Cd groups had significantly elevated brain Cd concentrations compared with controls (both *p* < 0.0001), whereas the AKK group alone did not differ from the control group (*p* = 0.998). Notably, the AKK+Cd group exhibited significantly higher brain Cd levels than the Cd-only group (*p* = 0.0236). These findings indicate that *A. muciniphila* treatment did not reduce brain Cd accumulation in mice; rather, it was associated with elevated brain Cd concentrations. The protective effects conferred by *A. muciniphila* against Cd-induced neurotoxicity therefore cannot be explained by reduced brain Cd accumulation but instead points to downstream mechanisms that preserve cognitive function despite an increased brain Cd burden.

### *A. muciniphila* normalized the Cd-induced dysregulation of hippocampal transcripts involved in learning and memory, vascular function, and oxidative stress in mice

Because *A. muciniphila* treatment preserved cognitive function without reducing brain Cd levels, we next investigated whether the treatment modulates downstream transcriptional responses in the hippocampus, the brain region most relevant to the NOL behavioral phenotype. Bulk RNA sequencing was performed to characterize transcriptional changes in the mouse hippocampus **(Fig. 4A and Supplementary Table 5)**. A total of 31 genes were differentially expressed between the Cd and AKK+Cd groups. Among them, 15 genes showed higher mRNA expression and 16 genes showed lower mRNA expression in the AKK+Cd group as compared to the Cd-only group **(Fig. 4B)**. Among the 15 genes with higher expression in the AKK+Cd group as compared to the Cd-only group, 8 were downregulated by Cd as compared to vehicle control and were restored by *A. muciniphila* treatment. These genes are *Klf8, Npas3, Penk, Psd4, Kcne2, Lypd1, Otof*, and Aqp1 **(Fig 5A)**. To further assess the functional relevance of these genes, we performed GO biological process and Mouse Genome Informatics (MGI) phenotype annotation analyses of representative learning and memory related genes. GO analysis revealed that *Penk*, *Otof*, and *Lypd1* were associated with neuropeptide signaling, cholinergic signaling, and synaptic vesicle and neurotransmitter release pathways **(Fig. 5B)**. In addition, MGI phenotype analysis indicated that *Lypd1*, *Npas3*, *Otof*, and *Penk* were linked to multiple phenotypes related to learning, memory, synaptic function, hippocampal morphology, and behavioral deficits **(Fig. 5C)**.

**Figure 4.**
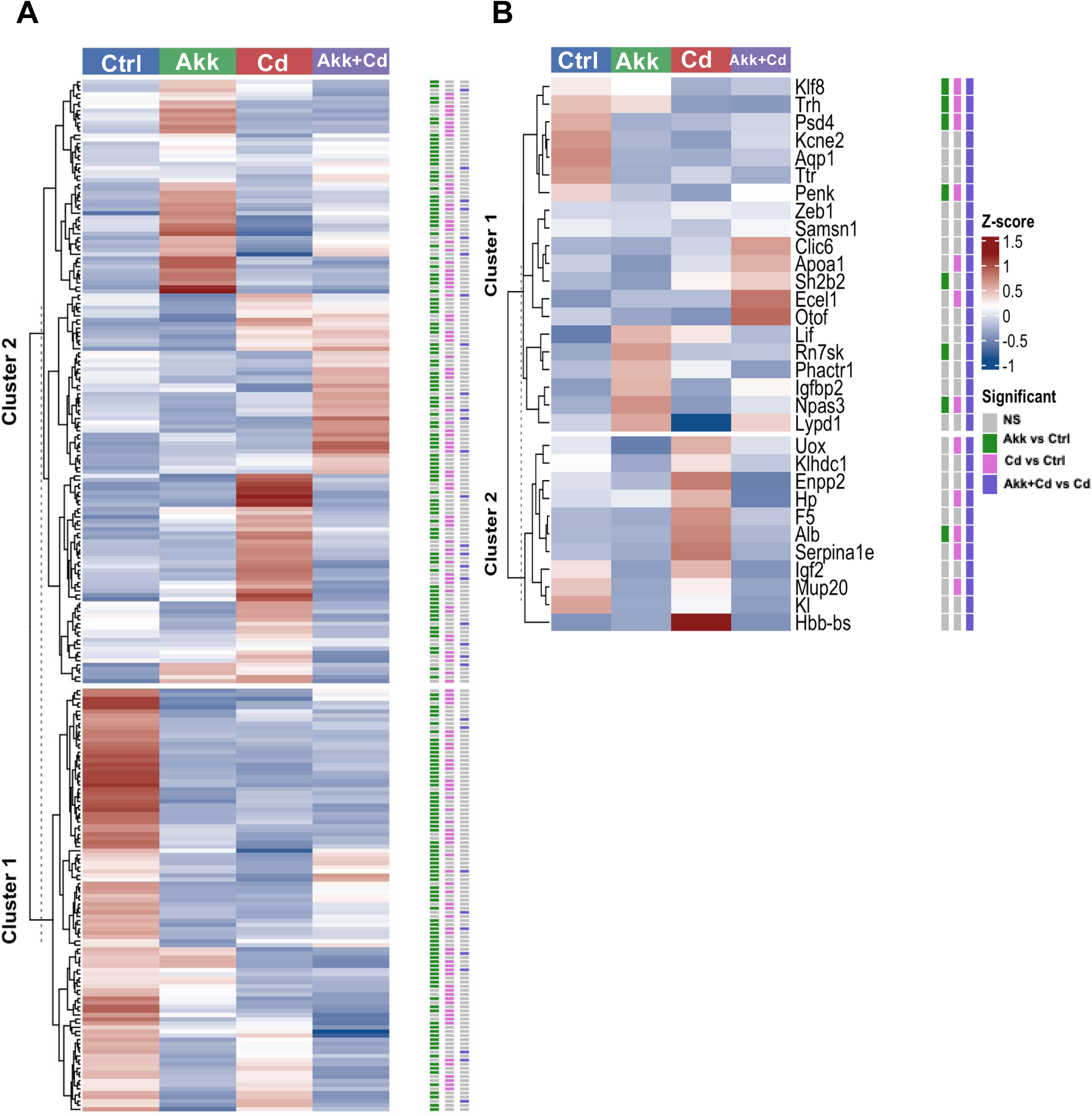
Hippocampal transcriptomic alterations induced by *A. muciniphila* treatment and Cd exposure. **A.** Heatmap showing hierarchical clustering of differentially expressed genes in the hippocampus across the four experimental groups (Ctrl, AKK, Cd, and AKK+Cd). **B.** Heatmap of the 31 differentially expressed genes identified between the Cd and AKK+Cd groups. Among these, 15 genes showed higher expression and 16 genes showed lower expression in the AKK+Cd group relative to the Cd group, indicating that A. muciniphila treatment reversed a subset of Cd-induced hippocampal transcriptional alterations. Color intensity indicates row-scaled expression levels (z-scores). The annotation bars on the right indicate whether each gene was significantly different in the AKK vs Ctrl, Cd vs Ctrl, or AKK+Cd vs Cd comparison. n = 4 in each group.

**Figure 5.**
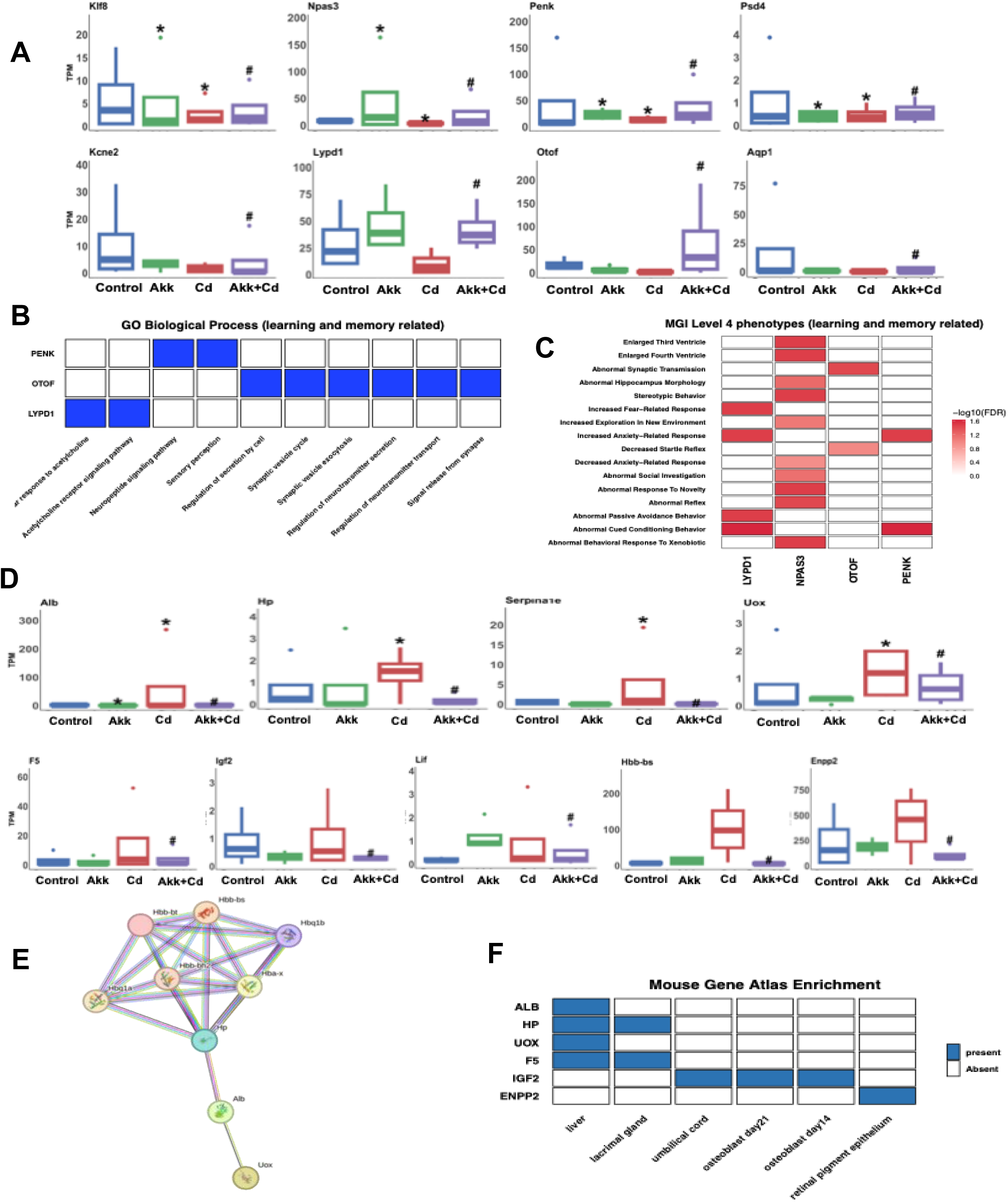
*A. muciniphila* reversed Cd-induced hippocampal transcriptional alterations related to neuronal function and vascular-associated stress signals. **A.** Box plots showing representative genes with lower expression in the Cd group and restored expression in the AKK+Cd group, including *Klf8, Npas3, Penk, Psd4, Kcne2, Lypd1, Otof,* and *Aqp1*. **B.** Gene Ontology (GO) biological process enrichment analysis of representative learning- and memory-related genes, highlighting enrichment in neuropeptide signaling, cholinergic signaling, and synaptic vesicle and neurotransmitter release pathways. **C.** Mouse Genome Informatics (MGI) phenotype enrichment analysis of representative learning- and memory-related genes, showing associations with phenotypes related to hippocampal morphology, synaptic transmission, and behavioral abnormalities. **D.** Box plots showing representative genes with higher expression in the Cd group and reduced expression in the AKK+Cd group, including *Alb, Hp, Serpina1e, Uox, F5, Igf2, Lif, Hbb-bs,* and *Enpp2*. **E.** STRING protein-protein interaction network of representative genes elevated in the Cd group and reduced in the AKK+Cd group, showing a closely connected network centered on hemoglobin-related genes and abundant plasma proteins. **F.** Mouse Gene Atlas enrichment analysis of representative Cd-elevated genes, showing enrichment in peripheral tissue categories, particularly liver-related signatures. n = 4 in each group. Data in **A** and **D** are presented as box plots showing the median, interquartile range, and individual data points. * *p* < 0.05 versus control; # *p* < 0.05 versus Cd.

Among the 16 genes that were downregulated in the AKK+Cd group as compared to the Cd-only group, 9 were up-regulated by Cd as compared to the control group, whereas *A. muciniphila* co-exposure attenuated the Cd effect. These genes include *Alb, Hp, Serpina1e, Uox, F5, Igf2, Lif, Hbb-bs,* and *Enpp2* **(Fig 5D)**. These genes are primarily associated with oxidative stress, vascular dysfunction, inflammatory responses, and metabolic disturbance. STRING protein-protein interaction analysis revealed a closely connected network centered on hemoglobin-related genes together with abundant plasma proteins including haptoglobin (*Hp*), albumin (*Alb*), and urate oxidase (*Uox*) **(Fig. 5E)**. Mouse Gene Atlas enrichment analysis further showed that several of these genes, including *Alb*, *Hp*, *Uox*, *F5*, *Igf2*, and *Enpp2*, were enriched in peripheral tissues, particularly liver-related categories **(Fig. 5F)**. Their presence in the hippocampus likely indicates blood-derived or vascular-associated signals, possibly resulting from residual blood components or altered blood-brain barrier integrity rather than intrinsic transcriptional changes within the brain.

Together, these findings demonstrate that *A. muciniphila* reversed Cd-induced hippocampal transcriptional dysregulation by restoring synaptic and learning- and memory-related genes, while suppressing Cd-elevated vascular, inflammatory, oxidative stress, and blood-derived transcriptional signatures. This pattern a resilience mechanism that operates downstream of brain Cd accumulation.

### Pre-treatment with *A. muciniphila* modified the effects of Cd exposure on the mouse gut microbiome and functional pathways

To investigate the gut-brain axis mechanisms underlying *A.muciniphila*-mediated cognitive protection, we examined how *A. muciniphila* treatment and Cd exposure affect the gut microbial community. To screen for microbial taxa showing treatment-dependent longitudinal changes, we performed longitudinal differential abundance analysis using linear mixed-effects models that accounted for repeated measurements within individual mice. For each taxon, abundance was modeled as a function of group, time, and the group x time interaction, with subject included as a random intercept. The global group x time interaction term was used to identify taxa whose abundance trajectories differed across the four experimental groups over the exposure period. Using a nominal significance threshold of *p* < 0.05, 65 candidate taxa with treatment-dependent temporal patterns during Cd exposure were identified for subsequent analyses **(Fig. 6A)**.

**Figure 6.**
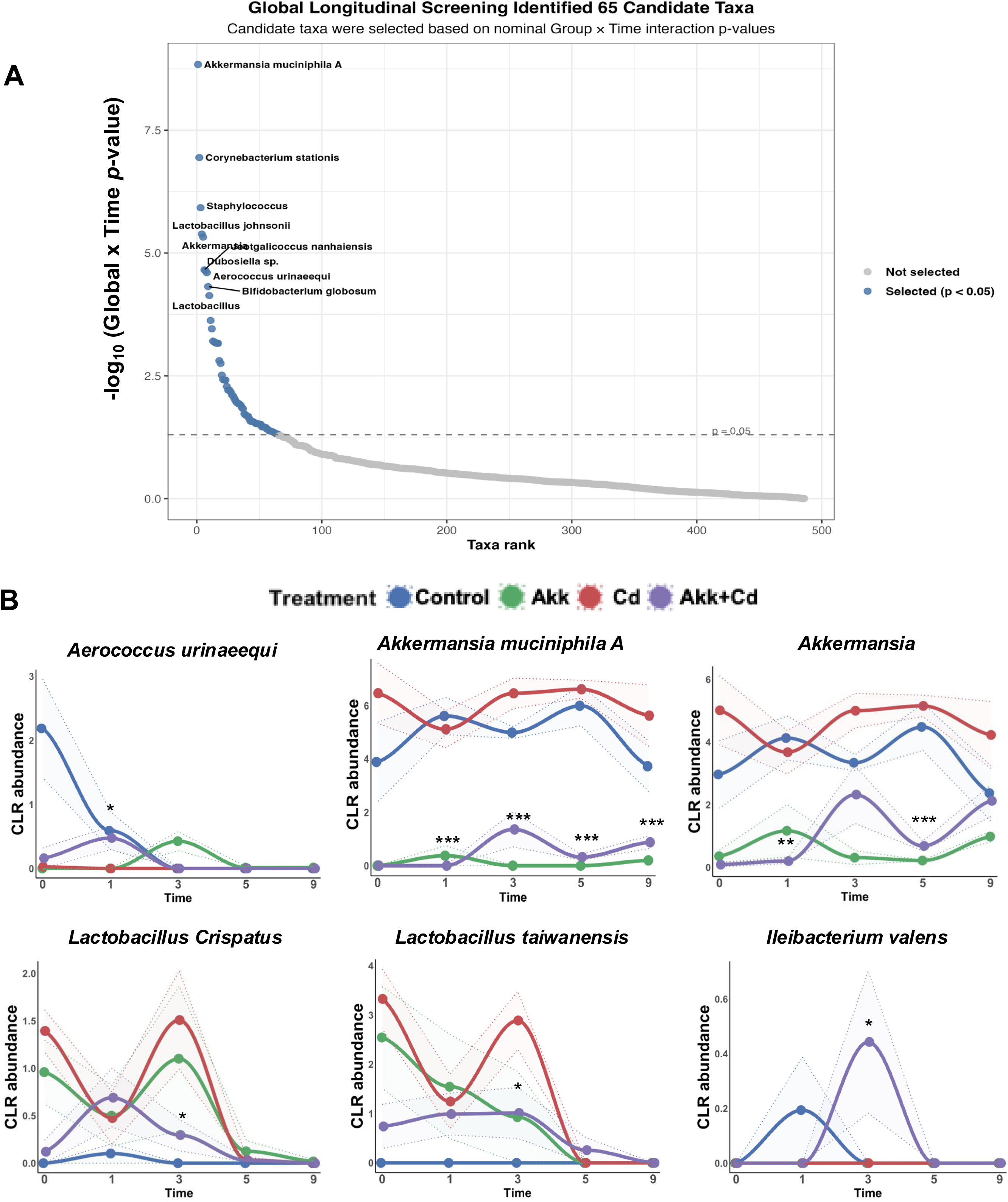
Candidate taxa screening and identification of gut microbial taxa differentially affected by *A. muciniphila* treatment during Cd exposure. **A.** Global longitudinal screening of gut microbial taxa based on nominal Group x Time interaction p-value from linear mixed-effects models. Taxa with nominally significant longitudinal differences (p < 0.05) were selected as candidate taxa for further analysis. Each point represents one taxon, ranked by the significance of the global Group × Time interaction. **B.** Longitudinal trajectories of CLR-transformed abundance for the six taxa showing significant week-specific differences between the Cd and AKK+Cd groups after Benjamini-Hochberg FDR correction. n = 4 in each group. Asterisks indicate significant week-specific differences between the Cd and AKK+Cd groups .* *q* < 0.05, ** *q* < 0.01, \*\*\**q* < 0.001.

For these 65 candidate taxa identified by the initial longitudinal screening, we next conducted targeted post hoc comparisons between the Cd and AKK+Cd groups during the active Cd exposure period (weeks 1, 3, 5, and 9). Baseline and week 0 samples were retained in the mixed-effects models to improve estimation of the overall longitudinal structure but were excluded from post-hoc comparisons. For each taxon, both the overall Group x Time interaction and week-specific Cd vs. AKK+Cd contrasts were evaluated **(Supplementary Table 3)**. Following Benjamini-Hochberg FDR correction, 6 taxa showed significant week-specific differences between the Cd and AKK+Cd groups, including *Aerococcus urinaeequi, Akkermansia muciniphila A*, *Akkermansia*, *Lactobacillus crispatus*, *Lactobacillus taiwanensis*, and *Ileibacterium valens* **(Fig. 6B)**. Notably, *A. muciniphila A* was consistently more abundant in the Cd group at weeks 1, 3, 5, and 9, and the genus *Akkermansia* was also enriched in the Cd group at weeks 1 and 5. This pattern contrasts with the increased *A. muciniphila* signal detected by qPCR **(Fig. 2B)** and the consistently positive log BAA-835:Strain 139 ratios observed in AKK-treated groups **(Fig. 2C)**. Our metagenomic profiling revealed that CLR abundance of endogenous Akkermansia-related taxa declined significantly in AKK-treated groups as early as one week after *Akkermansia* treatment, prior to Cd exposure **(Fig. S3)**, indicating that human-derived Akkermansia gavage itself suppressed resident murine Akkermansia populations. This suppression persisted through the exposure period for *A. muciniphila A*, while genus-level *Akkermansia* showed a trend toward recovery in the AKK+Cd group by week 9 (*p* = 0.058). Together, these observations suggest that BAA-835 colonization reshapes the resident murine *Akkermansia* community through niche remodeling rather than simple additive colonization. In addition, *L. crispatus* and *L. taiwanensis* were more abundant in the Cd group at week 3, whereas *I. valens* was increased in the AKK+Cd group at the same time. At week 1, *A. urinaeequi* was also more abundant in the AKK+Cd group. After FDR correction, no taxa showed a significant overall difference in longitudinal trajectory between the Cd and AKK+Cd groups. Collectively, these results demonstrate that *A. muciniphila* treatment selectively reshaped the temporal dynamics of specific gut microbial taxa during Cd exposure.

To further characterize the functional implications of these microbial compositional changes, we performed differential abundance analysis of predicted KEGG enzyme annotations at L2 and L3 levels, using the same analytical framework applied to the longitudinal taxa analysis **(Supplementary Table 4)**. At the L2 level, the most pronounced differences between the Cd and AKK+Cd groups emerged at week 5 **(Fig.7)**. Seven functional categories showed significant week-specific differences (FDR-adjusted *p* < 0.1): five were enriched in the AKK+Cd group, including Family S8: subtilisin family, Subtilisin family, AgrC-AgrA, Phosphate and amino acid transporters, and 5’ processing factors, while lipid transporters and Autotransporter (AT-1) family were significantly reduced. At the L3 level, a broader set of 20 specific pathways exhibited significant differences (FDR-adjusted *p* < 0.1). Among the 15 upregulated pathways in the AKK+Cd group, several were related to nutrient transport, including dipeptide transporter, putative glutamine transporter, cystine transporter, taurine transporter, AI-2 transporter, and phosphate transporter, as well as sugar-specific transporter systems such as Ascorbate-specific II component and Fructose specific II component. Pathways involved in proteolysis (Family S8: subtilisin family, Subtilisin family) and cell division regulation (Inhibitors of FtsZ assembly) were also enriched in the AKK+Cd group. In addition, five L3 pathways were downregulated in the AKK+Cd group, including the Autotransporter-1 (AT-1) family, lysophospholipid transporter (LplT) family, vitamin B12 transporter, transposing S-S bonds (5.3.4), and oxidoreductases using quinone or related compounds as acceptors (1.2.5).

**Figure 7.**
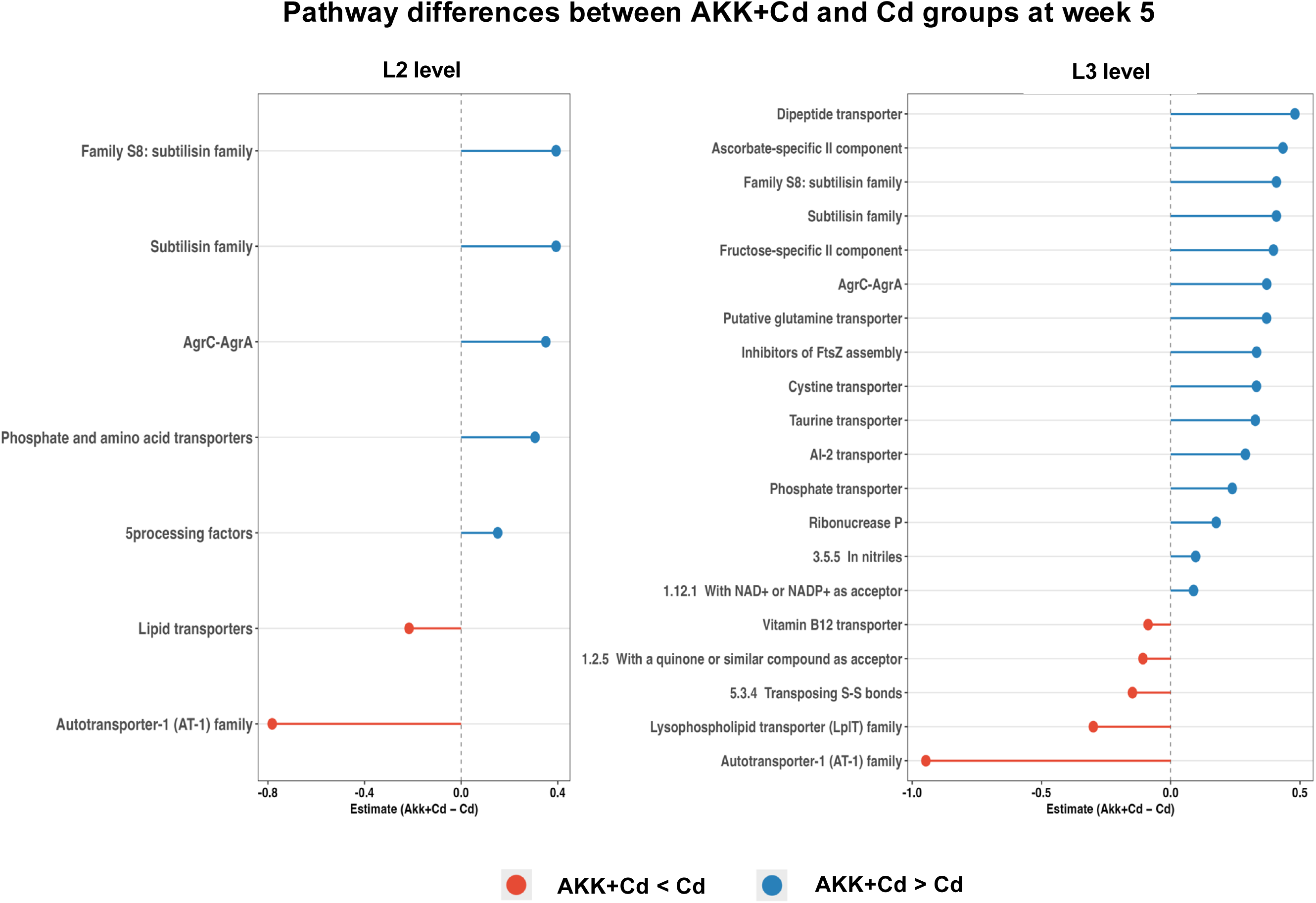
Gut microbial enzyme functions differentially altered between the AKK+Cd and Cd groups at week 5. Predicted gut microbial enzyme functions, annotated to the KEGG enzyme (EC) hierarchy, that were significantly altered between groups at week 5 by differential abundance analysis are shown at level 2 and level 3. Positive estimates indicate higher abundance in the AKK+Cd group, whereas negative estimates indicate lower abundance in the AKK+Cd group. n = 4 in each group.

To further examine the temporal dynamics of globally significant pathways, we traced week-specific AKK+Cd vs. Cd difference across weeks1, 3, 5, and 9 **(Fig. S4)**. These pathways were selected from the longitudinal screening step based on nominally significant global group x time interaction effects (*p* < 0.05). Notably, several pathways enriched in the AKK+Cd at week 5, such as AgrC-AgrA and phosphate and amino acid transporters, shifted toward depletion by week 9, suggesting a transient functional effect of *A. muciniphila* pre-treatment. In contrast, the AT-1 family remained consistently depleted across multiple time points. Together, these findings suggest that *A. muciniphila* pre-treatment reshaped the functional capacity of the gut microbiome during Cd exposure, particularly in nutrient transport, proteolysis, and membrane-associated pathways.

### *Lactobacillus* species are strongly associated with learning and memory performance in mice

Having identified taxa whose abundance differed between the Cd and AKK+Cd groups, we next asked which of these taxa are functionally relevant to cognitive performance. To investigate the potential correlation between the gut microbiome alterations and the learning and memory deficits, we selected a panel of high-confidence taxa for behavioral correlation analysis from the 21 taxa that showed week-specific differences at FDR-adjusted *p* < 0.1 between the Cd and AKK+Cd groups. Because cognitive deficits in NOL test emerged in the Cd group, but not in the AKK+Cd group, starting at week 5, we prioritized taxa showing significant differences between these two groups at weeks 3 and 5, as these were more likely to reflect biologically meaningful responses to Cd exposure and *A. muciniphila* treatment. We further excluded *Akkermansia*-related taxa to avoid treatment-driven circularity and to focus the analysis on endogenous downstream responders rather than the intervention organism itself. We further restricted the analysis to species-level taxa to improve precision. This refinement process yielded a final panel of eight high-confidence taxa for behavioral correlation analysis, including *Lactobacillu crispatus, Lactobacillu intestinalis, Lactobacillus taiwanensis, Paramuribaculum sp001689535, Ileibacterium valens, Bifidobacterium animalis, Faecalimonas umbilicate, and Flavonifractor sp000508885* **(Supplementary Table 3)**.

Repeated-measures correlation analysis was performed between CLR-transformed taxon abundance and the discrimination index (DI) from the NOL test results at weeks 3, 5, and 9 in the Cd and AKK+Cd groups. We found significant positive correlations for *L. crispatus* (r = 0.63, FDR-adjusted *p*= 0.0476*), L. intestinalis* (r = 057, FDR-adjusted *p* = 0.0476), and *L. taiwanensis* (r = 0.57, FDR-adjusted *p* = 0.0476) **(Fig.8)**. No significant correlations were detected for remaining five taxa. Because all three taxa showing significant positive correlations belong to the genus *Lactobacillus*, we further examined the correlation between *Lactobacillus* abundance and DI and discovered a significant positive correlation (r = 055, *p* = 0.0227) **(Fig. S5A)**. Together, these findings suggest that *Lactobacillus* taxa are positively associated with hippocampus-dependent learning and memory performance in mice.

**Figure 8.**
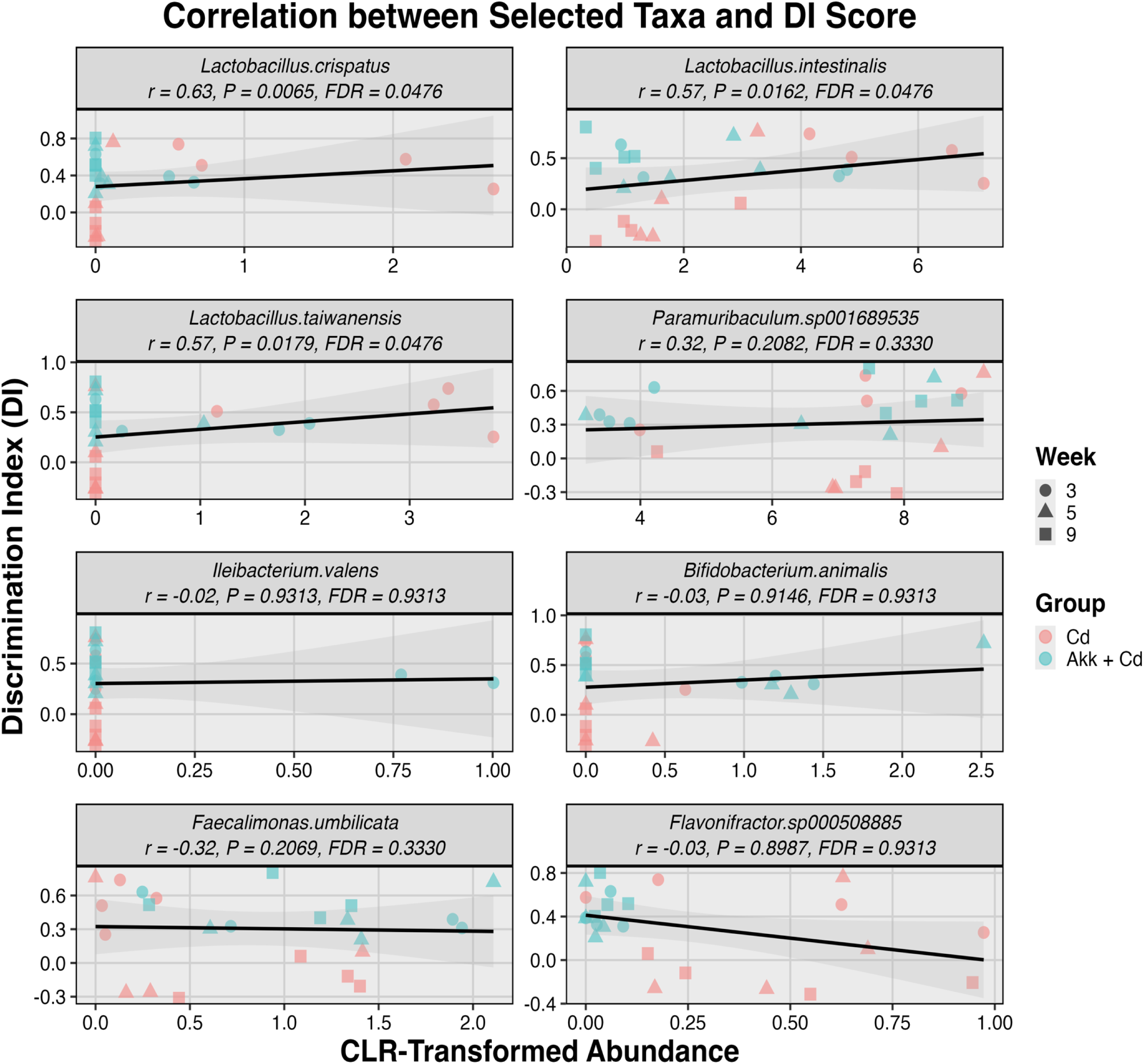
Correlation analysis identified associations between *Lactibacillus* taxa and spatial memory performance. Repeated-measures correlation analysis was performed between CLR-transformed abundance of selected taxa and DI values from the NOL test at weeks 3, 5, and 9 in the Cd and AKK+Cd groups. Points represent individual samples, with colors indicating treatment groups and shapes indicating time points. Black lines show repeated-measures correlation fits with 95% confidence intervals. n = 4 in each group.

### *A. muciniphila* treatment prevented the Cd-induced decrease in *Lactobacillus* taxa

Since *Lactobacillus* taxa showed significant positive associations with hippocampus-dependent spatial memory and differed significantly in abundance between the Cd and AKK+Cd groups, we further investigated the effects of *A. muciniphila* treatment on *Lactobacillus* at both the species and genus levels. Longitudinal abundance changes were modeled in the Cd and AKK+Cd groups across baseline, week 0 (one week after *A. muciniphila* treatment), and weeks 1, 3, 5, and 9 of Cd exposure. At the species level, *L. crispatus*, *L. intestinalis*, and *L. taiwanensis* all showed distinct temporal patterns between groups **(Fig. 9A)**. In the Cd group, all three taxa showed marked reductions relative to week 0, particularly at weeks 5 and 9. Interestingly, no significant decreases were detected in the AKK+Cd group, indicating that *A. muciniphila* treatment preserved these taxa during Cd exposure. Consistent with this pattern, week-specific comparisons of ΔCLR abundance (vs. week 0) **(Fig. 9B)** further showed significant differences for *L. crispatus* at week 1, 5, and 9. For *L. intestinalis*, significant differences were observed at week 1 and week 5, with a borderline difference observed at week 9 (FDR-adjusted *p* = 0.05). For *L. taiwanensis*, a significant difference was observed at week 5, with a borderline difference at week 9 (FDR-adjusted *p* = 0.05). At the genus level, we observed a similar protective effect of *A. muciniphila* treatment on *Lactobacillus* **(Fig. S5 B and C)**. Together, these findings demonstrate that *A. muciniphila* supplementation prevented the progressive Cd-induced decline in Lactobacillus taxa, consistent with a gut microbiome-mediated mechanism of cognitive protection.

**Figure 9.**
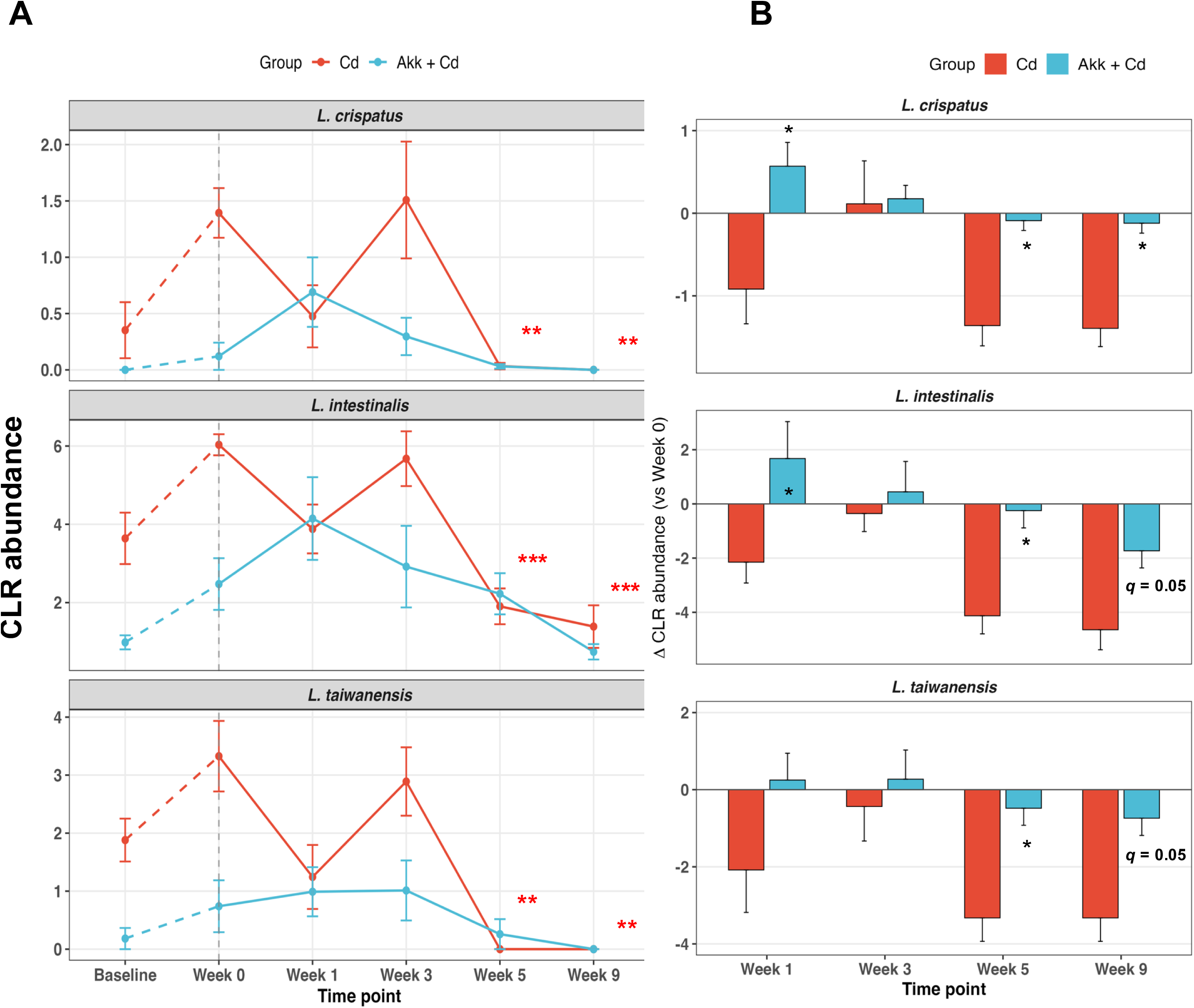
A. muciniphila treatment prevented the Cd-induced decline in *Lactobacillus* taxa. **A.** Longitudinal changes in CLR-transformed abundance of *Lactobacillus crispatus*, *Lactobacillus intestinalis*, and *Lactobacillus taiwanensis* in the Cd and AKK+Cd groups across baseline, week 0, and weeks 1, 3, 5, and 9 after Cd exposure. **B.** Changes in CLR abundance relative to week 0 (ΔCLR abundance) for the same three *Lactobacillus* taxa in the Cd and AKK+Cd groups at weeks 1, 3, 5, and 9. n = 4 in each group. Data are presented as mean ± SEM. Red asterisks in (A) indicate significant indicate significant within-group differences relative to week 0 in Cd group. Black asterisks in (B) indicate significant between-group differences at the indicated time points. * *q* < 0.05; ** *q* < 0.01; *** *q* < 0.001.

### *A. muciniphila* treatment attenuated Cd-impaired intestinal barrier integrity

The preservation of *Lactobacillus* taxa by *A. muciniphila* further indicated a gut-localized mechanism of protection. Since *Lactobacillus* species are known to support intestinal barrier integrity and our previous study demonstrated that Cd exposure impairs intestinal barrier integrity in mice ^21^, we next examined whether *A. muciniphila* treatment preserves barrier integrity under Cd exposure. We examined the mRNA expression of tight junction genes in the ileum and colon by RT-qPCR. In the ileum **(Fig. 10)**, Cd exposure caused a broader disruption of tight junction gene expression, with significant reductions in Claudin 1, 2, and 7(*Cldn1, Cldn2,* and *Cldn7*), Tight Junction Protein 1 and 2 (*Tjp1* and *Tjp2)*, Junctional Adhesion Molecule 1 and 2 (*Jam1* and *Jam2*) compared with the controls. Notably, *A. muciniphila* treatment significantly restored the expression of *Cldn7* and *Jam1* in the AKK+Cd group relative to the Cd group, while *Jam2* showed a trend toward recovery (*p* = 0.07). The expression of *Cldn1*, *Cldn2*, *Tjp1*, and *Tjp2* was also higher in the AKK+Cd group than in the Cd group, although these differences were not statistically significant. In the colon **(Fig. 11)**, Cd exposure significantly reduced the expression of *Cldn2*, *Jam1*, and *Tjp2* compared with the control group. *A. muciniphila* treatment restored the expression of these genes in the AKK+Cd group, although the differences relative to the Cd group did not reach statistical significance. In addition, *Cldn1* showed a trend toward recovery in the AKK+Cd group compared with the Cd group (*p* = 0.059). Together, these results indicate that Cd exposure disrupted intestinal barrier-related gene expression in both the ileum and colon, and that *A. muciniphila* treatment exerted a protective effect, with the most prominent effects observed in the ileum.

**Figure 10.**
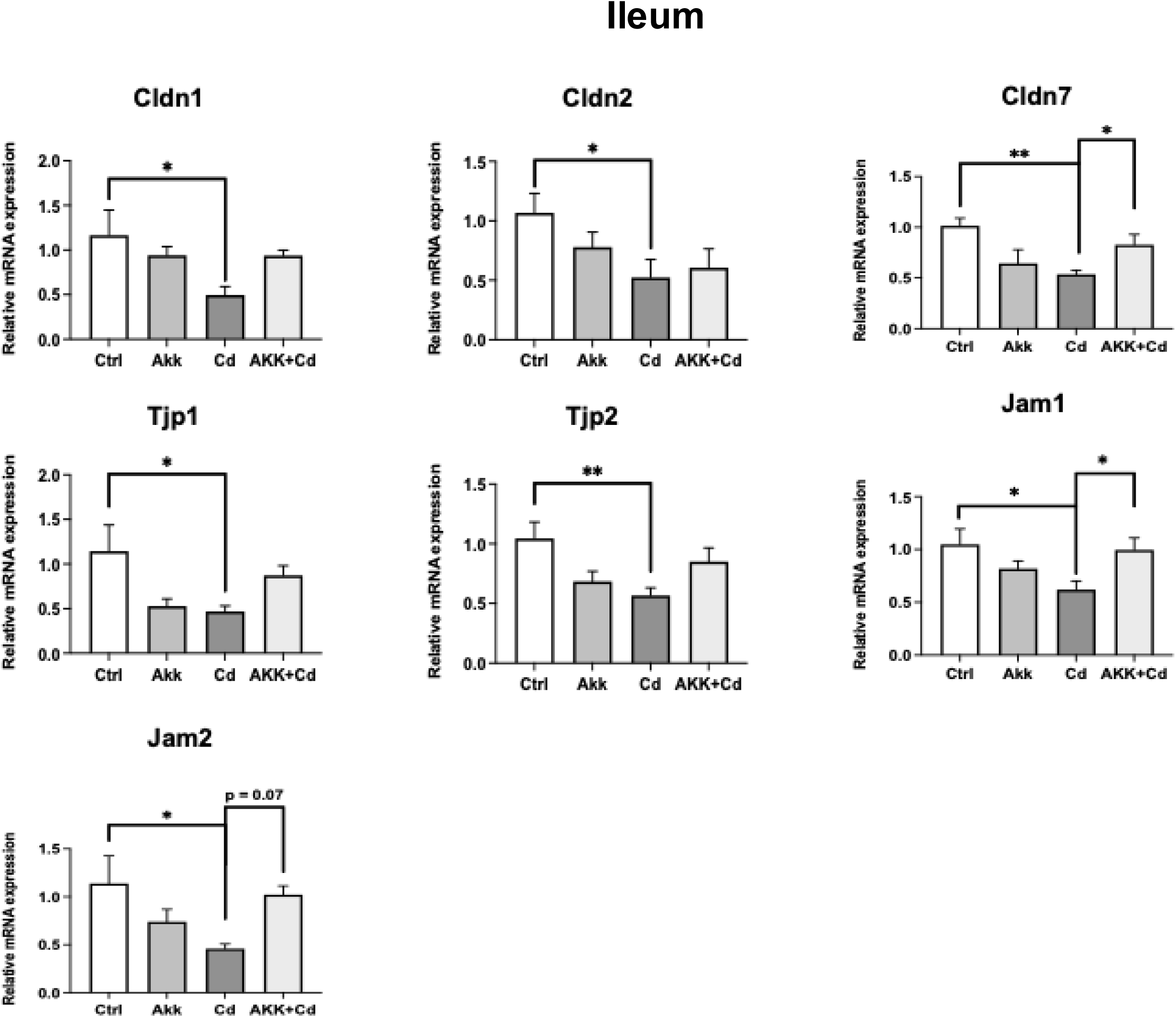
Effects of Cd exposure and *A. muciniphila* treatment on colon tight junction gene expression. Relative mRNA expression of tight junction-related genes was determined by RT-qPCR. Data are presented as mean ± SEM. n = 5-6 in each group. * *p* < 0.05; ** *p* < 0.01; *** *p* < 0.001.

**Figure 11.**
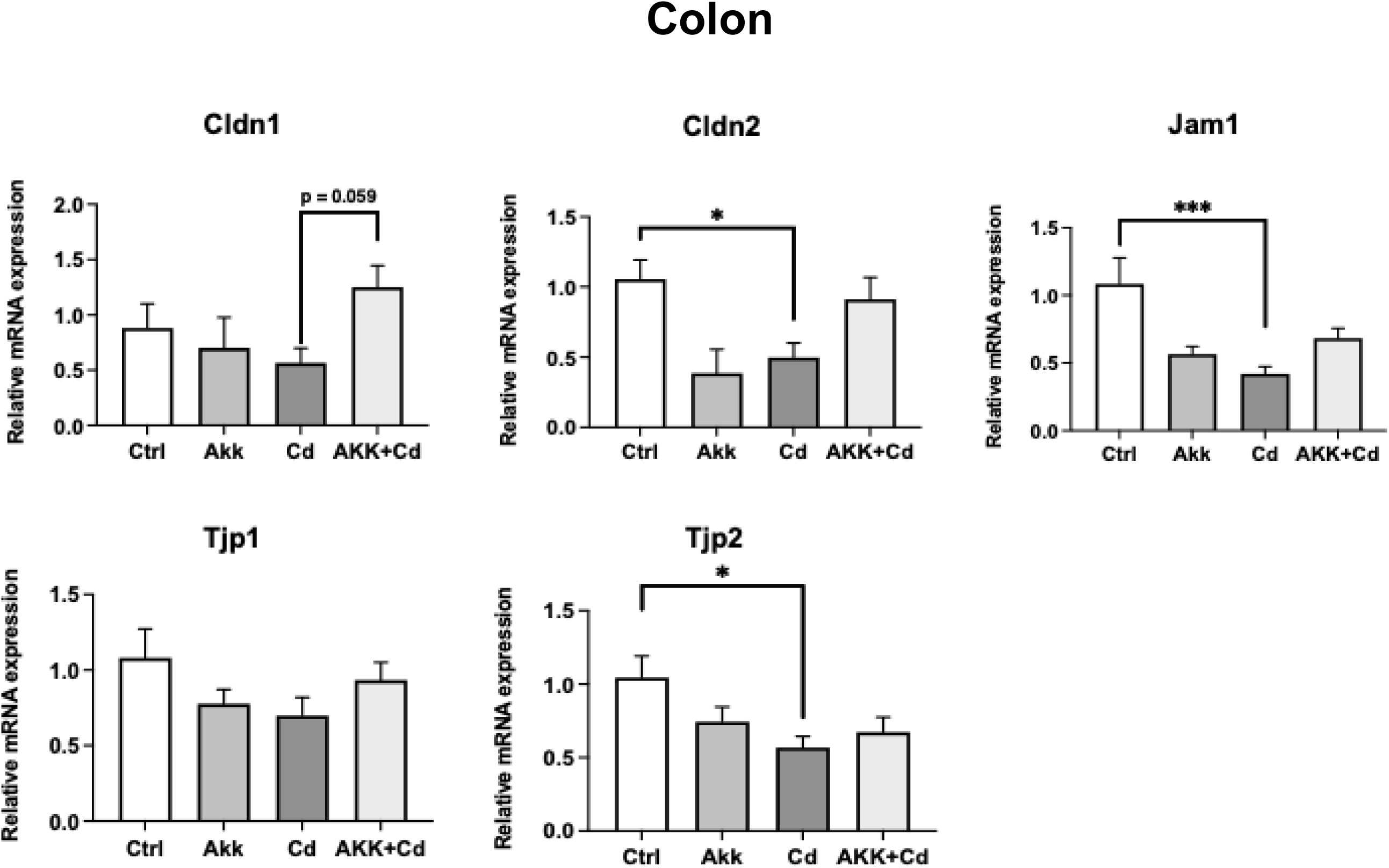
Effects of Cd exposure and *A. muciniphila* treatment on ileum tight junction gene expression. Relative mRNA expression of tight junction-related genes was determined by RT-qPCR. Data are presented as mean ± SEM. n = 5-6 in each group. * *p* < 0.05; ** *p* < 0.01; *** *p* < 0.001.

### *A. muciniphila* treatment attenuated Cd-induced serum cytokine dysregulation

Intestinal barrier disruption allows bacterial products and inflammatory mediators to enter the systemic circulation, potentially driving systemic inflammation that has been implicated in neuroinflammation and cognitive impairment ^21^. Given that *A. muciniphila* treatement protected the gut barrier integrity from Cd-induced disruption, we next quantified serum cytokine levels to determine the effects of Cd as well as the interactions between Cd and *A. muciniphila* on systemic inflammation. As shown in **Fig. 12**, serum Interleukin (IL)-1α and -10 levels were significantly elevated in the Cd group compared with the controls, and *A. muciniphila* co-exposure normalized both to levels comparable to controls. To note, While IL-1α is a pro-inflammatory cytokine ^54^, IL-10 is an anti-inflammatory cytokine that is commonly induced as a compensatory and regulatory response during inflammation ^55^. The concurrent Cd-induced elevation of both cytokines suggests a state of immune dysregulation characterized by simultaneous pro-inflammatory activation and compensatory anti-inflammatory signaling, a pattern that was reversed by A. muciniphila co-exposure.

**Figure 12.**
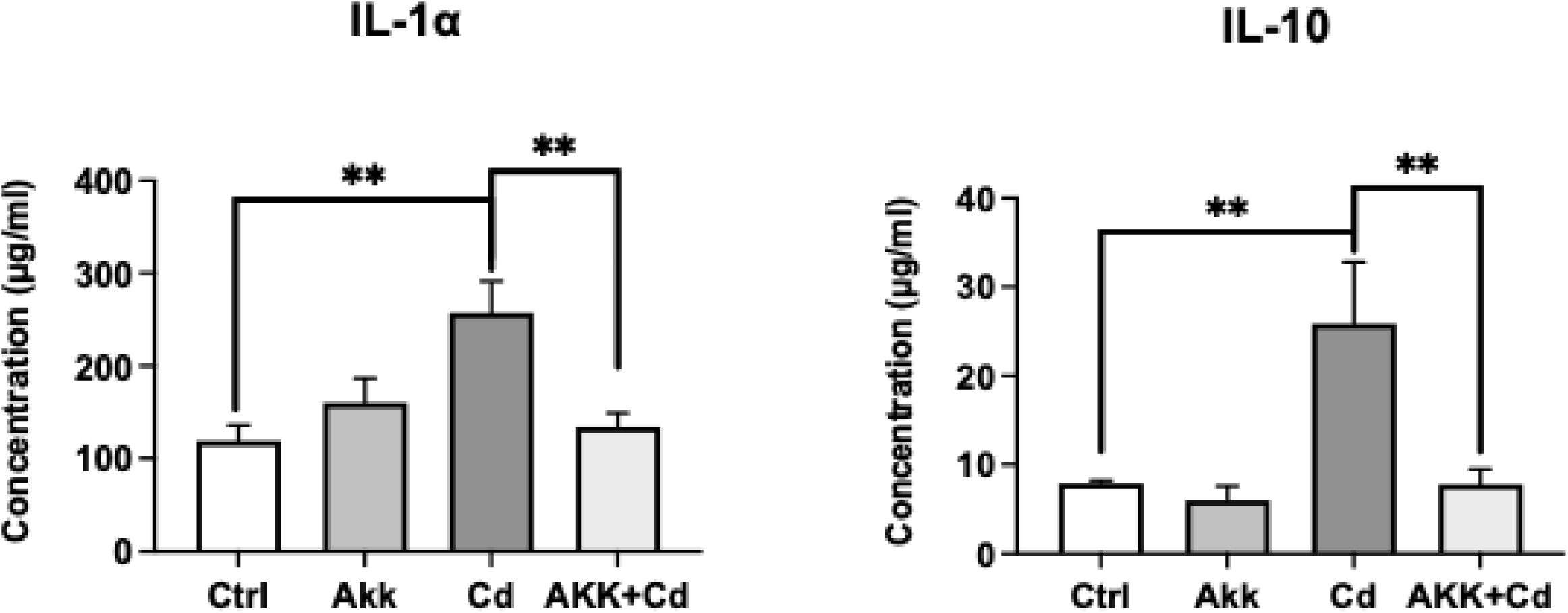
*A. muciniphila* treatment attenuated Cd-induced alterations in serum inflammatory cytokines. Data are presented as mean ± SEM. n = 4-5 in each group. * *p* < 0.05; ** *p* < 0.01; *** *p* < 0.001.

### *A. muciniphila* treatment modulated gut short-chain fatty acid profiles altered by Cd exposure

In addition to physical barrier maintenance and inflammatory regulation, microbial metabolites are another key mediators through which the gut microbiome communicates with the brain ^14^. Short-chain fatty acids (SCFAs) are downstream metabolites of *A. muciniphila* and other gut commensals, and many of them are known to regulate intestinal barrier integrity, inflammation, and learning and memory ^13,14^. To determine whether *A. muciniphila*-mediated cognitive protection against Cd neurotoxicity involves changes in microbial metabolites, we quantified SCFAs in large intestine content (LIC), small intestine content (SIC), whole brain, and serum by GC-MS. Interestingly, *A. muciniphila* had limited effects on most SCFAs detected, with the most notable directional shift in SIC acetic acid. Cd exposure increased SIC acetic acid to 1.7-fold that of controls (Cd = 30.9 ± 7.8 vs. Ctrl = 18.1.6 ± 5.7 µg/mL), whereas *A. muciniphila* co-exposure resulted in the lowest acetic acid concentrations among all groups (Akk+Cd = 8.6 ± 2.9 µg/mL), representing a 72% reduction relative to Cd-only group. Two-way ANOVA showed a trend-level AKK main effect (*p* = 0.06), with no significant Cd main effect or Cd x AKK interaction; Tukey HSD comparison AKK+Cd vs. Cd approached but did not reach conventional significance (*p* = 0.096). In the LIC, 2-methylpentanoic acid, a branched-chain fatty acid (BCFA) derived from bacterial protein fermentation, showed a strong *A. muciniphila*-associated effect among all detected SCFAs. Cd exposure numerically elevated LIC 2-methylpentanoic acid (Cd = 1.18 ± 0.05 vs. Ctrl = 1.02 ± 0.04 µg/g), whereas two *A. muciniphila*-supplemented groups showed comparable reductions (AKK = 0.95 ± 0.05; AKK+Cd = 0.95 ± 0.04 µg/g). Two-way ANOVA revealed a significant *A. muciniphila* main effect (*p* = 0.043), indicating that *A. muciniphila* consistently lowered LIC 2-methylpentanoic acid regardless of Cd exposure. Tukey HSD comparisons showed directional reductions in both AKK vs. Cd (*p* = 0.115) and AKK+Cd vs. Cd (*p* = 0.120), consistent with the overall *A. muciniphila* main effect. **(supplemental Figs 6-9)**.

To examine whether *A. muciniphila*-responsive SCFAs changes were associated with cognitive function, Spearman correlation analysis was performed between the NOL DI and pre-specified metabolites. Correlation analysis was restricted to metabolites showing AKK-related changes, defined as a significant AKK main effect (*p* < 0.05) or a Cd x AKK interaction (*p* < 0.05) Based on these criteria, six metabolites were selected for correlation analysis: isovaleric acid, 2-methylbutyric acid, 2-methylpentanoic acid, caproic acid, and heptanoic acid in LIC, and serum caproic acid. As shown in **Figs. 13 and 14**, LIC 2-methylpentanoic acid showed a significant negative correlation with DI (ρ = -0.714, *p* = 0.0019, FDR-adjusted *p* = 0.011), indicating that higher colonic 2-methylpentanoic acid levels were associated with poorer cognition. Two additional SCFAs correlations, LIC 2-methylbutyric acid (ρ = -0.508, *p* = 0.045, FDR-adjusted *p* = 0.089) and LIC isovaleric acid (ρ = -0.511, *p* = 0.043, FDR-adjusted *p* = 0.089), were nominally significant but did not survive FDR correction. Notably, all these correlations were negative and involved BCFAs, whereas none of the straight-chain SCFAs tested showed significant correlations with DI. Consistent with these findings, fecal isovaleric acid has been reported to be negatively correlated with cognition in AD patients ^56^. Together, these findings identify BCFAs, particularly colonic 2-methylpentanoic acid, rather than straight-chain SCFAs, as the metabolites most closely linked to Cd-induced cognitive decline, and as candidate gut-derived mediators of *A. muciniphila* protection.

**Figure 13.**
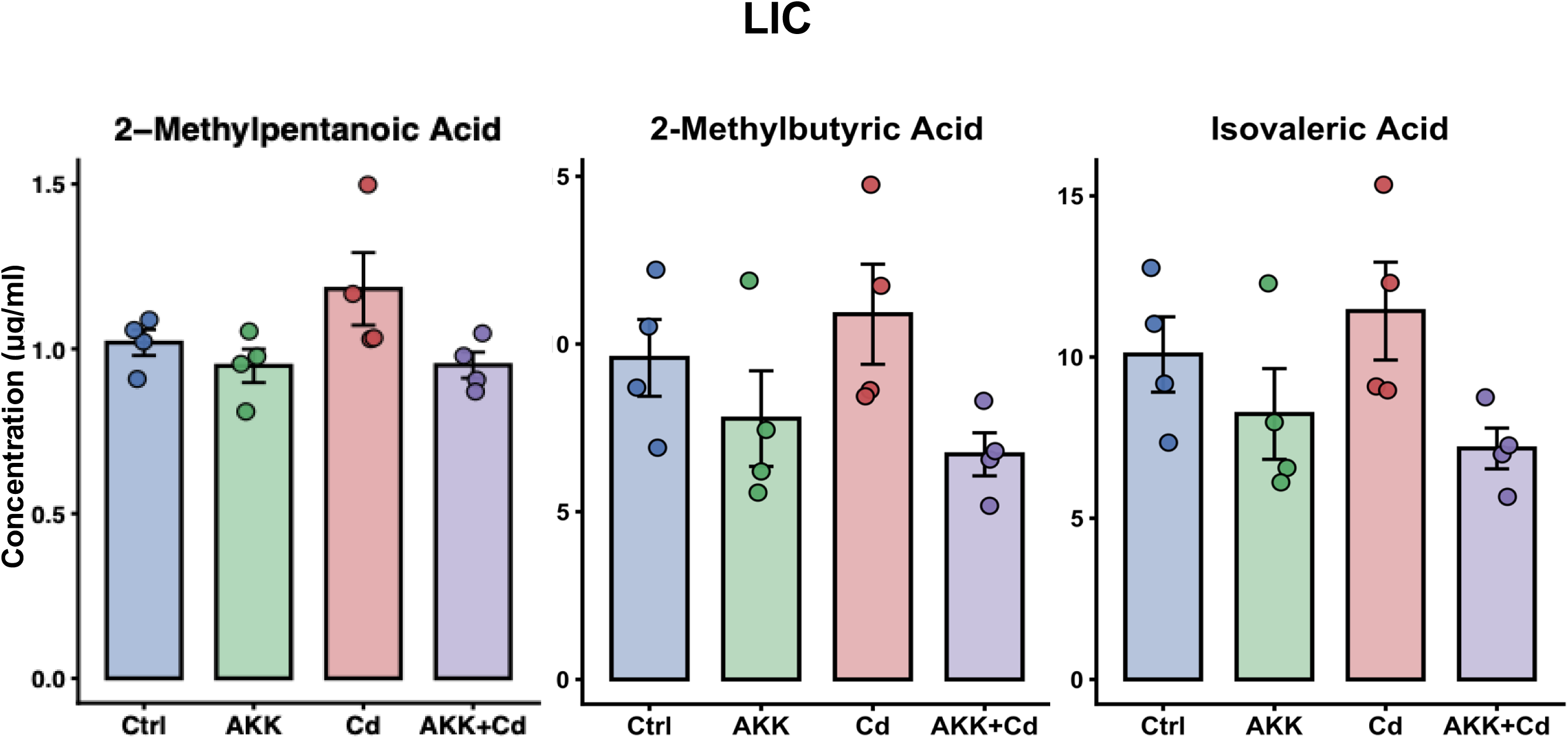
Concentrations of 2-methylpentanoic acid, 2-methylbutyric acid and isovaleric acid in LIC. Data are presented as mean ± SEM. n = 3-4 in each group.

**Figure 14.**
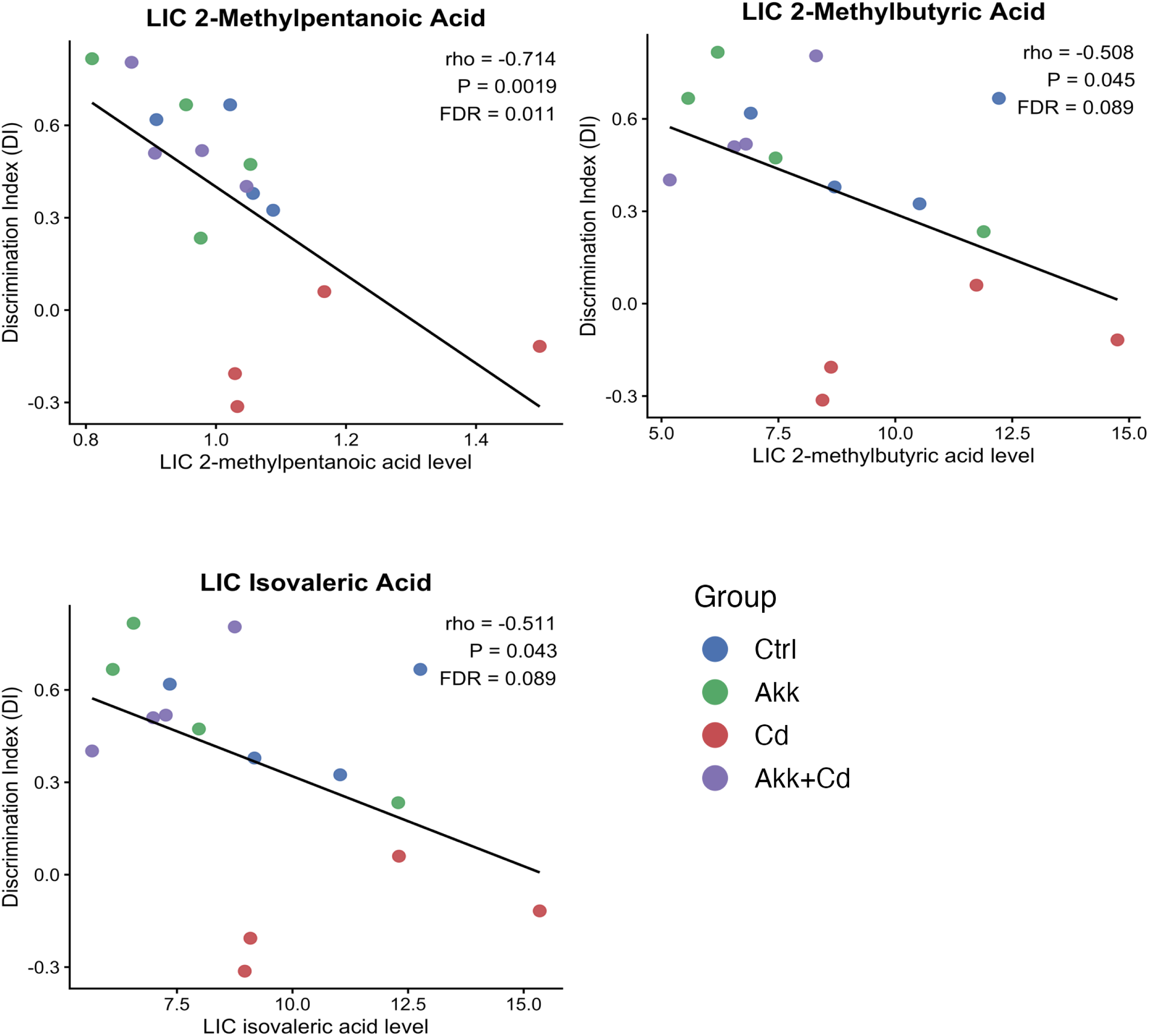
LIC 2-methylpentanoic acid, 2-methylbutyric acid and isovaleric acid are negatively correlated with spatial memory performance. Spearman correlation analyses were performed between the discrimination index (DI) and four selected metabolites. n = 4 in each group. Reported statistics include Spearman’s ρ, nominal *p* values, and FDR-adjusted *p* values.

## Discussion

In the present study, we demonstrate that *A. muciniphila* supplementation prevents Cd-induced hippocampus-dependent cognitive impairment through a mechanism independent of reduced brain Cd accumulation. This protective effect was accompanied by coordinated changes across the gut-brain axis, including preservation of *Lactobacillus* taxa, restoration of intestinal barrier-related gene expression, attenuation of systemic inflammatory dysregulation, modulation of microbial metabolites, and reversal of Cd-associated transcriptional alterations in hippocampal genes involved in synaptic and vascular functions. Notably, these benefits occurred despite elevated brain Cd concentrations in the AKK+Cd group, indicating that the protective effect of *A. muciniphila* operates independently of limiting Cd entry into the brain and instead engages downstream gut-brain signaling to preserve cognitive function even under conditions of increased brain metal burden. Our findings support a model in which gut microbial homeostasis can confer cognitive resilience against environmental neurotoxicant exposure independently of the brain toxicant distribution.

The protective effect observed in our study is notable because it was preventive rather than remedial. *A. muciniphila* was administered prior to Cd exposure and continued throughout the study, and the AKK+Cd group remained comparable to unexposed controls in the NOL test performance across the entire testing period **(Fig. 3A-D)**. This contrasts with most probiotic strategies reported for environmental neurotoxicity, which have generally been tested in treatment or co-administration paradigms and have shown variable efficacy ^36,37,57^. In our model, Cd-induced impairment developed progressively, first emerging at week 5 and maintaining significance throughout the study, suggesting that *A. muciniphila* prevented the onset of cognitive deficit rather than merely mitigating or reversing an established phenotype. This distinction has potential translational relevance: preventive microbiome-based strategies may be particularly well suited for populations chronically exposed to low-level Cd, such as individuals living in industrial or agriculturally contaminated environments, where early intervention could limit the gradual emergence of neurobehavioral impairment. Interestingly, *A. muciniphila* supplementation did not reduce brain Cd burden under Cd exposure **(Fig. 3E)**. Our data suggest that *A. muciniphila* preserves cognition not by preventing Cd accumulation in the brain, but by buffering its downstream molecular effects within the central nervous system and along the gut-brain axis, a mechanism we term as “tolerance through resilience”. The “tolerance through resilience” interpretation is supported by the hippocampal transcriptomic changes observed in the AKK+Cd group. Among the genes suppressed by Cd exposure and restored by *A. muciniphila*, a prominent subset is closely associated with neuronal function and cognition **(Fig.5)**. For example, KLF8 (Krüppel-like factor 8) is a transcription factor (TF) involved in the activation of the Wnt/β-catenin signaling pathway ^58^ ^59^. In the context of Alzheimer’s disease (AD), *Klf8* expression is reduced in patient brains, and increased *Klf8* expression has been shown to ameliorate AD-like pathology through Wnt/β-catenin signaling pathway activation in rodent AD model ^60,61^. Neuronal PAS domain protein 3 (NPAS3) is a hippocampus-expressed TF required for hippocampal neurogenesis ^62,63^. In mice, *Npas3* deficiency has been associated with behavioral abnormalities, including autism-like behaviors^64^. *Penk* encodes proenkephalin, the precursor of endogenous opioid peptides that modulate neurogenesis, synaptic transmission, and hippocampal plasticity, learning, and memory ^65,66^. The restoration of *Penk* expression by *Akkermansia* treatment therefore suggests a potential microbiome-dependent mechanism for preserving hippocampal neuropeptide signaling. In addition, *Lypd1* and *Otof* are functionally linked to cholinergic signaling, synaptic vesicle exocytosis, and neurotransmitter release ^67,68^ Together, restoration of the expression of these genes suggests that *A. muciniphila* preserves hippocampal transcriptional programs regulating the neuronal and synaptic function that were disrupted by Cd exposure.

Importantly, *A. muciniphila* also suppressed a distinct group of Cd-elevated hippocampal genes associated with vascular, inflammatory, and blood-derived signals **(Fig. 5)**. Several of these genes, including *Alb*, *Hp*, *Hbb-bs*, and *F5*, encode proteins typically confined to the circulating compartment. Their elevation in the hippocampus suggests possible extravasation following blood-brain barrier (BBB) compromise or expression by infiltrating immune cells following barrier disruption ^68–71^. For example, albumin (ALB) is a major plasma protein whose abnormal presence in brain tissue is widely considered a marker of BBB leakage ^68^, whereas F5 is a coagulation-related factor associated with vascular injury, inflammatory activation, and disruption of the neurovascular environment ^70^. This gene-expression pattern is consistent with previous reports that Cd exposure can impair BBB integrity ^72,73^, and that *A. muciniphila* can support barrier functions along the gut-brain axis ^41,74^. In addition, *Serpina1e* belongs to the murine Serpina1 family, which encodes alpha-1 antitrypsin–like serine protease inhibitors predominantly produced in the liver and circulate in the blood. Alpha-1 antitrypsin family proteins are closely linked to systemic inflammatory responses and vascular injury ^75^. In our study, the upregulation of *Serpona1e* under Cd exposure likely reflects activation of vascular or perivascular cells, or infiltration of peripheral immune cells following BBB disruption. *Uox*, another gene typically restricted to peripheral tissues, encodes urate oxidase, a liver-enriched enzyme involved in uric acid metabolism that is minimally expressed in the healthy mouse brain ^76^. Because urate metabolism is tightly connected to oxidative stress and redox homeostasis, elevated *Uox* expression in the hippocampus is more suggestive of peripheral metabolic or vascular-associated disturbances than of a neuron-specific response. Together, the reduction of these genes’ expression in AKK+Cd mice supports the hypothesis that *A. muciniphila* attenuates Cd-induced vascular, inflammatory, and oxidative stress–related signals in the hippocampal microenvironment. Direct validation at the protein, cellular, and blood-brain barrier levels can be conducted in future studies.

The gut microbiome data further support the gut-centered mechanism underlying the *A. muciniphila-*mediated protection. Rather than broadly reshaping the overall microbial community, *A. muciniphila* selectively modulated a subset of microbial taxa during Cd exposure. Among these, species in the *Lactobacillus* genus emerged as the most behaviorally relevant microbial responders **(Fig. 8)**. *Lactobacillus* taxa, including *L. crispatus, L. intestinalis, and L. taiwanensis*, were positively associated with spatial memory performance in the NOL test, and *A. muciniphila* administration effectively prevented their Cd-induced depletion **(Fig. 9)**. To note, *L. crispatus* and *L. taiwanensis* are also found in human fecal/gut microbiota ^77,78^, whereas *L. intestinalis* is better supported as an intestinal *Lactobacillus* originally isolated from rodent intestine ^79^. Importantly, because of natural inter-individual variation at baseline, the absolute abundance of *Lactobacillus* in the AKK+Cd cohort was not consistently higher than that in the Cd-only group at every time point. Rather, the Cd group exhibited a progressive and significant decline relative to baseline, whereas *A. muciniphila* supplementation completely prevented this downward trend.

The preservation of *Lactobacillus* taxa may represent one downstream consequence of this ecological stabilization. Previous studies have consistently shown that *Lactobacillus* species contribute to gut-brain communication and can induce beneficial effects on cognitive function through modulation of intestinal barrier function, neuroimmune responses, microbial metabolites, and hippocampal plasticity-related pathways ^80,81^. Several *Lactobacillus* strains, though from species different from those detected in our study, have been reported to improve cognitive deficits in models of environmental neurotoxicants or systemic stress. For instance, *L. rhamnosus* GR-1 attenuated lead-induced spatial learning and memory deficits ^82^, whereas *L. rhamnosus* GG improved sepsis-related cognitive impairment ^83^. *L. johnsonii* BS15 prevented stress-induced memory dysfunction, further supporting a role for *Lactobacillus* in maintaining cognitive function ^84,85^. Among the three *Lactobacillus* taxa identified in our study that were positively associated with hippocampus-dependent learning and memory, *L. intestinalis* has been linked most directly to gut-brain communication. Prior work has shown that *L. intestinalis* can influence host behavior through vagus nerve-dependent signaling ^86^, and that it suppresses intestinal inflammation, modulates mucosal immune responses, and preserves epithelial homeostasis under conditions of intestinal injury or toxicant exposure ^87,88^. *L. intestinalis* has been reported to increase in parallel with improved spatial memory in a murine model of chemically induced cognitive impairment, though this evidence remains associative ^89^. In addition, *L. crispatus* is supported as a barrier- and immune-supporting taxon, with studies showing that it can improve epithelial integrity, promote epithelial repair, and modulate inflammatory responses, suggesting a potentially indirect contribution to cognitive resilience through preservation of mucosal homeostasis ^90,91^. Evidence for *L. taiwanensis* remains more associative, with its enrichment reported in parallel with improved spatial learning and memory in an aging mouse model, alongside potential antibacterial and intestinal immune-regulatory properties ^92,93^. Collectively, these observations support the broader premise that *Lactobacillus* taxa can contribute to cognitive resilience via the microbiota-gut-brain axis, although direct functional evidence specifically for *L. crispatus*, *L. intestinalis*, and *L. taiwanensis* in cognitive contexts remains limited and warrants dedicated investigation.

We also find that *A. muciniphila* prevents the decline of *Lactobacillus* taxa under Cd exposure. Notably, *A. muciniphila* administration alone did not induce a marked increase in *Lactobacillus* abundance, arguing against a simple growth-promoting effect. Instead, its impact became evident specifically under Cd exposure, where it effectively prevented the progressive loss of *Lactobacillus* species over time. Although direct species-to-species interactions between *A. muciniphila* and *Lactobacillus* remain to be established, previous studies provide a biologically plausible framework for such an effect. As a mucin-degrading bacterium, *A. muciniphila* can utilize host-derived mucin and generate metabolites that support other mucus-associated or saccharolytic bacteria, including certain *Lactobacillus* species ^94^. In addition, *A. muciniphila* has been widely reported to enhance mucosal barrier function, promote mucus-layer integrity, and modulate intestinal inflammation ^95–97^. In the context of Cd-induced toxicity, these observations suggest that *A. muciniphila* may function as an ecological buffer, stabilizing the gut microbial community structure and mucosal environment under external stress. Under this framework, the preservation of *L. crispatus*, *L. intestinalis*, and *L. taiwanensis* observed in our study may reflect secondary stabilization of microbial niches rather than direct promotion of these taxa.

The concurrent enrichment of proteolytic capacity (subtilisin family) and amino-acid/peptide transporters in the AKK+Cd group, alongside a trend toward lower concentrations of certain colonic BCFAs, may initially appear counterintuitive, since BCFAs are products of microbial branched-chain amino acid fermentation. One possible explanation is that enhanced bacterial uptake of amino-acid and peptide following *A. muciniphila* supplementation likely allows proteolysis products to be channeled preferentially into biomass synthesis rather than dissipative deamination-based fermentation. This metabolic redirection effectively reduces BCFA efflux into the lumen - a mechanism consistent with the broader concept of nitrogen-conserving microbial metabolism in saccharolytic-leaning ecosystems. In parallel, the downregulation of autotransporter-1 systems, known to drive adhesion, virulence, and epithelial damage in various Gram-negative pathogens ^98,99^, suggests a transition toward a less pathogenic functional profile in the *A. muciniphila*-supplemented community. This shift perfectly aligns with the concurrent restoration of intestinal barrier gene expression. Collectively, these pathway-level alterations indicate that *A. muciniphila* supplementation does not merely alter taxonomic composition but fundamentally reshapes gut microbial function toward a more nutrient -efficient and less pro-inflammatory metabolic state.

In addition to microbial compositional and functional changes, *A. muciniphila* also modulated key functional aspects of the gut, including barrier integrity, inflammatory response, and microbial metabolism. Disruption of intestinal barrier function is a key feature of toxicant-induced gut dysfunction and can facilitate translocation of proinflammatory signals into systemic circulation. Previous studies have demonstrated that *A. muciniphila* can enhance intestinal barrier integrity, promote tight junction assembly, and improve gut permeability, partly through tight-junction-related mechanisms ^31,100^. Consistent with this, Cd exposure decreased expression of tight junction genes, whereas *A. muciniphila* supplementation partially restored these alterations, particularly in the ileum **(Fig. 10 and 11)**. In addition, intestinal barrier disruption is frequently accompanied by elevated systemic inflammation **(Fig. 12)**, which has been closely linked to impaired hippocampal function and cognitive decline. *A. muciniphila* has been reported to exert anti-inflammatory effects, including reduction of proinflammatory cytokines and modulation of mucosal immune responses in various disease models ^47,101^. Consistent with this, in our study, Cd exposure altered serum cytokine profiles, increasing pro-inflammatory cytokine IL-1α and anti-inflammatory cytokine IL-10. The simultaneous increase in IL-1α and IL-10 suggests a state of immune dysregulation characterized by concurrent pro-inflammatory activation and compensatory anti-inflammatory signaling. Notably, *A. muciniphila* supplementation normalized these alterations, significantly reducing both IL-1α and IL-10 levels toward pre-exposure levels, suggesting that *A. muciniphila* mitigates Cd-induced immune dysregulation and helps restore immune homeostasis, rather than exerting a simple unidirectional anti-inflammatory effect.

Microbial metabolites serve as another critical mediator for gut-brain communication. Among these, short-chain fatty acids (SCFAs) are among the most extensively characterized, known to influence neural function through multiple mechanisms, including microglial maturation, BBB integrity, neurotransmitter biosynthesis, and vagal signaling ^102,103^. In our study, Cd exposure produced compartment-specific disturbances in SCFAs homeostasis, and *A. muciniphila* supplementation partially normalized these changes, with the clearest behavioral association observed for branched-chain fatty acids (BCFAs) **(Fig. 13 and 14)**. Cd exposure elevated acetic acid in the SIC, which was markedly reduced in AKK+Cd mice **(Fig. S7)**. This observed pattern parallels the restoration of ileal tight-junction gene expression **(Fig. 11)**, consistent with previous findings that Cd-induced disruption of small intestinal epithelial homeostasis, including impaired barrier integrity, contributes to luminal acetate accumulation, whereas *A. muciniphila* re-establishes intestinal metabolic balance by restoring these functions. However, neither small intestinal nor colonic acetic acid correlated with the behavioral results, indicating that the acetate response did not track individual variation in cognitive performance in our model. Prior studies suggest that microbiota-derived acetate can influence learning and memory through gut-brain pathways involving microglial maturation, neuroimmune regulation, and neural signaling ^104–106^. The acetate response observed in our study may therefore primarily reflect recovery of gut epithelial and metabolic homeostasis, while remaining biologically relevant to cognition through indirect mechanisms not fully captured by a simple metabolite-behavior correlation test.

In contrast, BCFAs showed the clearest behavioral associations. Colonic 2-methylpentanoic acid was strongly and negatively associated with hippocampus-dependent learning and memory, and additional BCFA-related metabolites showed trends in the same direction, whereas no straight-chain SCFA correlated with cognitive performance **(Fig. 14)**. BCFAs are produced through microbial deamination of branched-chain amino acids and have been proposed as markers of a pro-inflammatory, proteolytic gut environment ^107–109^. Elevated fecal and circulating BCFAs have been reported in gut dysbiosis contexts including aging ^110^ and high-protein/low-fiber dietary intake ^111,112^. Emerging evidence further links BCFAs specifically to cognitive deficits. For instance, serum BCFAs were implicated in working-memory deficits through IL-6- and IL-8-driven neuroinflammation in patients with bipolar disorder ^113^. In addition, elevated circulating BCAA metabolites, which serve as upstream substrates for microbial BCFA production, have been associated with AD and cognitive decline, with dietary BCAA restriction ameliorating AD-related pathology and cognitive deficits in mice ^114,115^. Together, this BCFA-specific pattern suggests that BCFA accumulation is the metabolite signature most proximally associated with Cd-induced cognitive deficits in this model. However, the precise mechanisms by which colonic BCFAs reach or influence the brain function are unknown and will require direct mechanistic investigation.

Taken together, the downstream changes observed in this study, including preservation of *Lactobacillus* taxa, restoration of intestinal barrier gene expression, attenuation of systemic immune dysregulation, constraint of colonic BCFA accumulation, and reversal of Cd-associated hippocampal transcriptional changes, support a coordinated rather than pathway-specific mechanism of protection. *A. muciniphila* appears to act not through a single mediator, but by stabilizing the broader gut ecosystem during Cd exposure, with downstream effects on barrier integrity, immune signaling, and microbial metabolite signaling profiles converging to preserve cognition. This “multi-pathway gut-brain resilience” framework may help explain why cognitive preservation remained robust despite elevated brain Cd concentrations. These findings are highlighted by comparison with recent work in a colitis-related cognitive deficits study. Chen et al. reported that orally administered *A. muciniphila*-derived extracellular vesicles (AmEVs) reversed colitis-related cognitive impairment through tryptophan metabolic reprogramming of the gut-brain axis ^41^. Consistent with our findings, they discovered that AmEVs treatment restored intestinal barrier integrity, normalized inflammatory signaling, and recovered hippocampal synaptic genes expression. The overlap in these downstream outcomes across two very different disease contexts suggests that *A. muciniphila* may protect cognition through a shared set of gut-brain mechanisms, even when the upstream triggers and specific metabolic mediators differ. Of note, ATCC BAA-835 used in our study was originally isolated from healthy human feces ^29^ and has been used in many preclinical studies ^57,116^, strengthening the translational relevance of the gut-brain protective mechanisms identified here to human populations chronically exposed to environmental toxicants.

Several limitations of the current study warrant consideration. First, while the longitudinal behavioral and microbiome analyses were well-powered, the SCFA and cytokine analyses were conducted at a single terminal time point using relatively small sample sizes. Future studies utilizing larger cohorts and multiple time points will be critical to capture the dynamic shifts of these mediators throughout the entire exposure period. Second, hippocampal transcriptomic analyses were performed on bulk tissue, limiting our ability to resolve cell-type-specific responses. Subsequent investigations employing single-cell RNA-sequencing would help dissect the relative contributions of neurons, microglia, astrocytes, endothelial cells, and infiltrating immune cells to the observed transcriptional alterations. Third, BBB perturbations were inferred from blood-associated transcriptional signatures rather than quantified directly; therefore, dedicated tracer- or imaging-based assays are needed in the future studies to explicitly assess BBB permeability. In addition, although *Lactobacillus* taxa and colonic BCFAs emerged as compelling candidate mediators of cognitive performance, their precise causal roles must still be established via targeted supplementation, depletion, direct metabolite administration, or gnotobiotic models. Furthermore, because this study employed a single Cd dose and a fixed *A. muciniphila* dosing regimen, evaluating dose-response relationships, optimal intervention timing, and the necessity of continuous versus finite supplementation remains an important future direction. This study was also conducted in male mice; given the potential sex differences in Cd neurotoxicity, gut microbiome composition, immune responses, and other important factors, future studies including both sexes will be needed to determine whether the protective effects of *A. muciniphila* generalize across sexes. Finally, the paradoxical elevation of brain Cd in the AKK+Cd group represents an unexpected finding whose mechanistic basis warrants dedicated future investigation.

In summary, our findings demonstrate that *A. muciniphila* supplementation prevents Cd-induced hippocampus-dependent learning and memory impairment through a mechanism independent of brain Cd accumulation **(Fig. 15)**. Rather than reducing brain Cd concentrations, *A. muciniphila* appears to stabilize gut microbial and host physiological responses to chronic Cd stress by preserving *Lactobacillus* populations, maintaining intestinal barrier integrity, normalizing systemic immune dysregulation, constraining proteolytic microbial fermentation, and safeguarding hippocampal transcriptional programs related to neuronal and vascular function. These findings identify *A. muciniphila* as a promising candidate for preventive microbiome-based intervention against environmental Cd neurotoxicity and support a broader “tolerance through resilience” concept for understanding how gut-resident probiotics may protect against neurotoxicants whose tissue distribution cannot be fully controlled.

**Figure 15.**
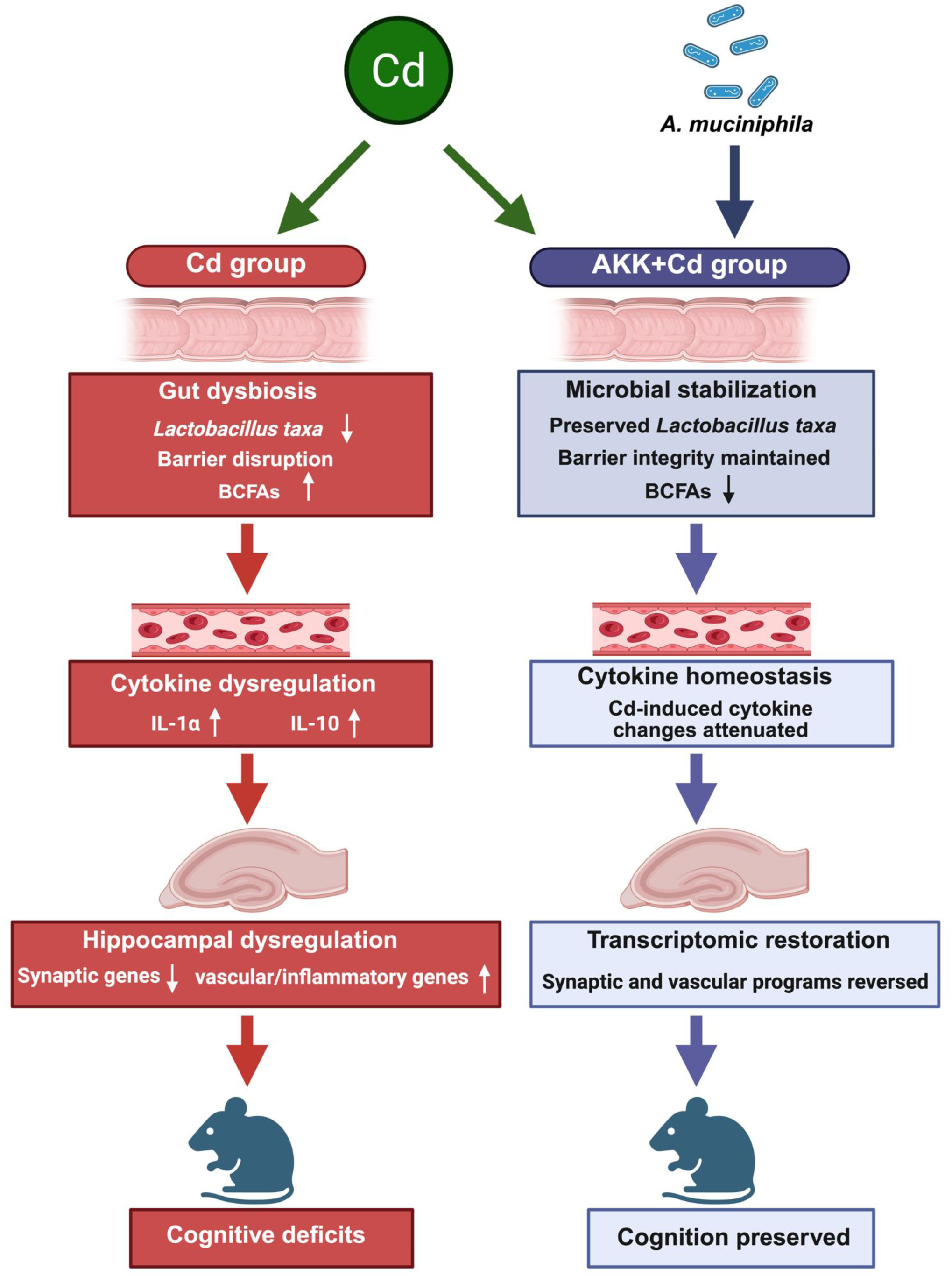
Summary of proposed mechanisms of *A. muciniphila*-mediated protection against Cd-induced cognitive impairments in mice.

## Supporting information

Supplemental figures with legends

## Acknowledgments

This research was supported by the National Institute of Environmental Health Sciences [R00ES034068 (to Dr. Wang); University of Michigan Lifestage Environmental Exposures and Disease Center (NIH P30ES017885); R01ES025708, R01ES030197, R01ES031098 (to Dr. Cui)]; National Institute of Diabetes and Digestive and Kidney Disease F31DK139707 (to Dr. Lim); National Institute of Aging U01AG088407; University of Washington Environmental Health and Microbiome Research Center (EHMBRACE), as well as University of Washington Center for Exposures, Diseases, Genomics, and Environment (NIH P30ES0007033). Oregon State University startup funding (to Dr. Cui); and the University of Washington Microbial Interactions and Microbiome Center (mim_c) award (to Dr. Lim). The authors would also like to thank the members of Dr. Wang’s and Dr. Cui’s laboratories for their help in tissue collection and manuscript revision.

## CRediT authorship contribution statement

**Hao Wang:** Conceptualization, Methodology, Investigation, Validation, Formal analysis, Data Curation, Visualization, Funding acquisition, Resource, Supervision, Writing – original draft. **Joe Jongpyo Lim:** Formal analysis, Visualization, Funding acquisition, Writing – Review & Editing. **Jinhua Chi:** Investigation, Data Curation. **Haiwei Gu:** Investigation, Writing – Review & Editing. **Julia Yue Cui:** Conceptualization, Methodology, Supervision, Funding acquisition, Resources, Writing – Review & Editing.

## Data availability statement

The metagenomic shotgun sequencing data generated in this study has been deposited in the NCBI Sequence Read Archive under accession number PRJNA1492592. The hippocampal RNA-seq data have been deposited in the NCBI Gene Expression Omnibus under accession number GSE338204.All other data supporting the findings of this study, including longitudinal microbiome and correlation analysis, RNA processed data, SCFAs data, qPCR primer sequences and results are provided in the supplementary materials. Additional information is available upon reasonable request.

## Declaration of Generative AI and AI-assisted Technologies

During manuscript preparation, the authors used ChatGPT (OpenAI, GPT-5.5) and Claude (Anthropic, Opus 4.7 and 4.8) to assist with English language editing, readability improvement, and drafting of data analysis code (e.g., R scripts for statistical analyses). These tools were not used to generate original scientific data, conduct final statistical analyses without author verification, or draw scientific conclusions. All AI-assisted outputs were critically reviewed, independently verified, and revised by the authors, who take full responsibility for the accuracy and integrity of the manuscript.

## Declaration of competing interest

The authors declare that they have no known competing financial interests or personal relationship that could have appeared to influence the work reported in this paper.

## Notes

### Competing Interest Statement

The authors have declared no competing interest.

