## Supplemental figures with legends for "*Akkermansia muciniphila* prevents cadmium-induced cognitive impairment through gut microbiome–mediated mechanisms": Supplementary figure legends (2).docx

**Figure S1. Strain-specific metagenomic mapping of A. muciniphila across the four treatment groups.** Fecal metagenomes from all treatment groups were aligned to the human-derived *A. muciniphila* (ATCC BAA-835, left panel) and the murine-derived (Strain 139, right panel) using Bowtie2. Time −1 represents baseline prior to any treatment; Time 0 represents one week after *A. muciniphila* gavage; Time 1, 3, 5, and 9 represent weeks following Cd exposure. n = 4 in each group.

**Figure S2. Open filed test results before and after treatment.** Open field test parameters were measured in control, AKK, Cd, and AKK+Cd mice before treatment and after the experimental intervention. No significant group differences were observed in locomotor activity- or anxiety-related measures, including movement time, travel distance, center exploration, margin exploration, and speed. n = 9-10 in each group. Data are shown as mean ± SEM.

**Figure S3. Longitudinal trajectories and baseline-to-week0 changes in Akkermansia-related taxa. A.** CLR-transformed abundance of *Akkermansia muciniphila A*, *Akkermansia muciniphila*, and genus-level *Akkermansia* across the study period. Baseline represents samples collected before *A. muciniphila* administration, and week 0 represents samples collected after one week of *A. muciniphila* administration and before Cd exposure. * at week 0 indicate within-group comparisons versus baseline. # at week 9 indicates endpoint comparisons between AKK+Cd and Cd groups. † at week 9 indicates endpoint comparisons between Akk and Control groups. **B.** Baseline-to-week 0 changes in CLR abundance, calculated as week 0 minus baseline for each mouse. Points represent individual mice, boxed show the median and interquartile range, and the dashed line indicates no change from baseline. * indicates comparisons of each treatment group versus Controls. All comparisons were performed using estimated marginal means from linear mixed-effects models for panel A and linear models for panel B, with Benjamini-Hochberg-adjusted p values. * *p* < 0.05, ** *p* < 0.01, ****p* <0.001.

**Figure S4. Temporal dynamics of globally significant gut microbial functional pathways in the AKK+Cd and Cd groups.** Bubble plots showing week-specific differences in CLR-transformed pathway abundance between the Akk+Cd and Cd groups at weeks 1, 3, 5, and 9. Pathways shown were selected from the longitudinal screening step based on nominally significant global Group × Time interaction effects (*p* < 0.05). The left panel shows level 2 (L2) functional categories, and the right panel shows level 3 (L3) categories. Blue circles indicate pathways enriched in the AKK+Cd group relative to the Cd group, whereas red circles indicate pathways depleted in the AKK+Cd group. n = 4 in each group

**Figure S5. Genus-level association and longitudinal dynamics of *Lactobacillus* during Cd exposure. A.** Repeated-measures correlation between CLR-transformed *Lactobacillus* abundance and DI in the Cd and Akk+Cd groups at weeks 3, 5, and 9. **B.** Longitudinal abundance trajectories of the *Lactobacillus* genus across baseline, week 0, and post-Cd exposure time points. (C) ΔCLR abundance relative to week 0, showing that the Cd-induced decline in *Lactobacillus* abundance was attenuated in the Akk+Cd group. Red asterisks in (A) indicate significant indicate significant within-group differences relative to week 0 in Cd group. Black asterisks in (B) indicate significant between-group differences at the indicated time points. n = 4 in each group. * *p* < 0.05; ** *p* < 0.01; *** *p* < 0.001.

**Figure S6. Short-chain fatty acids in serum**. Data are presented as mean ± SEM. n = 3-4 in each group. * p < 0.05; ** p < 0.01; *** p < 0.001.

**Figure S7. Short-chain fatty acids in small intestinal content**. Data are presented as mean ± SEM. n = 3-4 in each group. * p < 0.05; ** p < 0.01; *** p < 0.001.

**Figure S8. Short-chain fatty acids in large intestinal content**. Data are presented as mean ± SEM. n = 3-4 in each group. * p < 0.05; ** p < 0.01; *** p < 0.001.

**Figure S9. Short-chain fatty acids in brain**. Data are presented as mean ± SEM. n = 3-4 in each group. * p < 0.05; ** p < 0.01; *** p < 0.001.

**Table S1. Primer sequences used in RT-qPCR**

| **Target Genes** | **Forward Primer Sequence** | **Reverse Primer Sequence** |
| --- | --- | --- |
| *Cldn1* | GGACTGTGGATGTCCTGCGTTT | GCCAATTACCATCAAGGCTCGG |
| *Cldn2* | AGGACTTCCTGCTGACATCCAG | AATCCTGGCAGAACACGGTGCA |
| *Cldn7* | CTGCCTTGGTAGCATGTTCCTG | CCAGCCGATAAAGATGGCAGGT |
| *F11r (Jam1)* | CACCTACTCTGGCTTCTCCTCT | TGCCACTGGATGAGAAGGTGAC |
| *Jam2* | CAGACTGGAGTGGAAGAAGGTG | GCTGACTTCACAGCGATACTCTC |
| *Tjp1* | TTAAGCCTCCGGAAGTAGCA | GGAGCCTGTAGAGCGTTTTG |
| *Tjp2* | AATGGAAAGGTTGGCAACTG | CGTGCTTGTCCTGCTCAATA |
| *Gapdh* | GGCAAATTCAACGGCACAGT | GTCTCGCTCCTGGAAGATGG |
| *β-actin* | GGCCAACCGTGAAAAGATGA | CAGCCTGGATGGCTACGTACA |
| *Akkermansia muciniphila* | TAGTCCAGGGCATGCAGAAA | ATCCCTGAAAACAACCGTGC |
