## Supplemental figures with legends for "*Akkermansia muciniphila* prevents cadmium-induced cognitive impairment through gut microbiome–mediated mechanisms": Supplementary Figure_v9.pdf

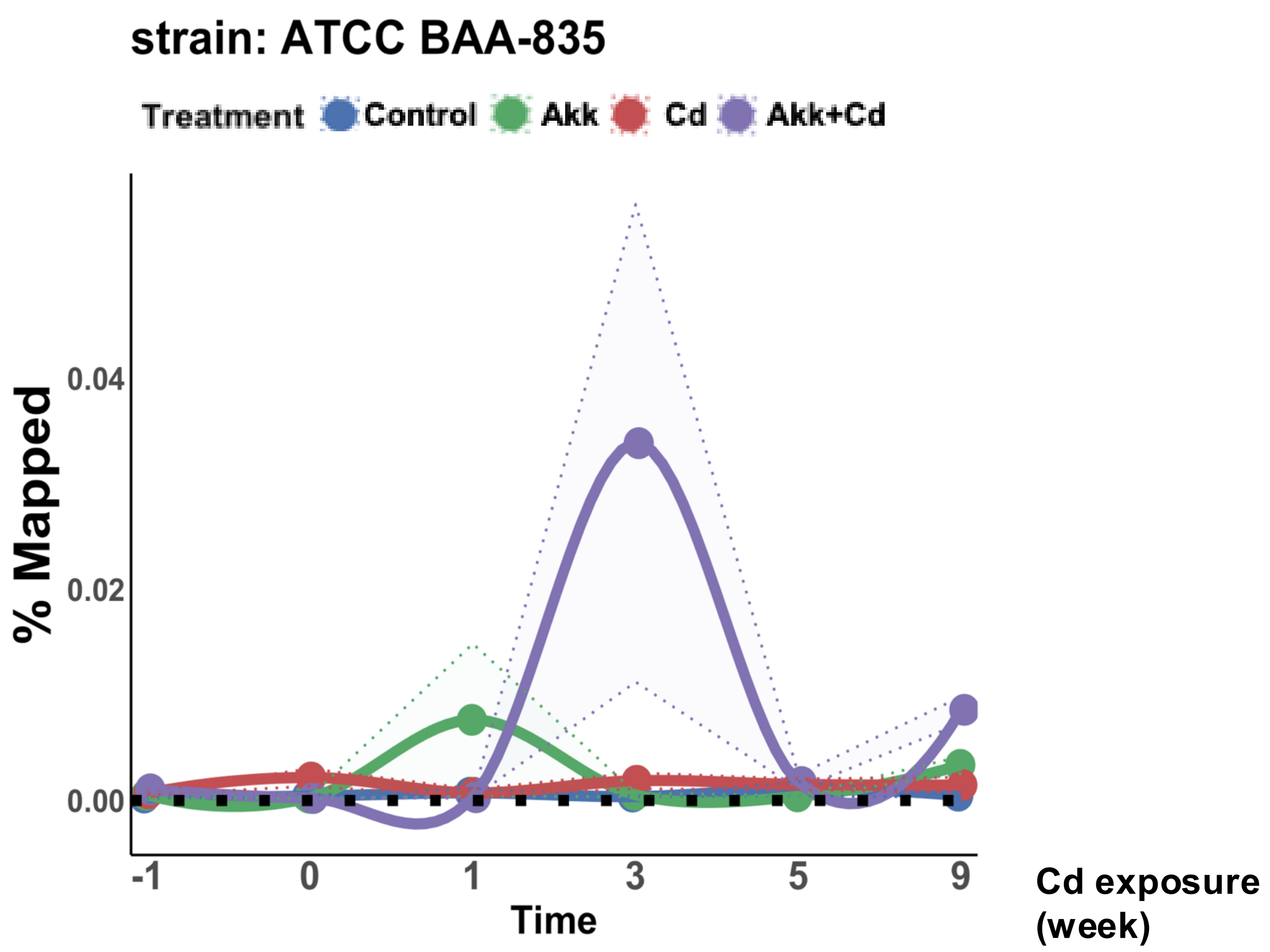


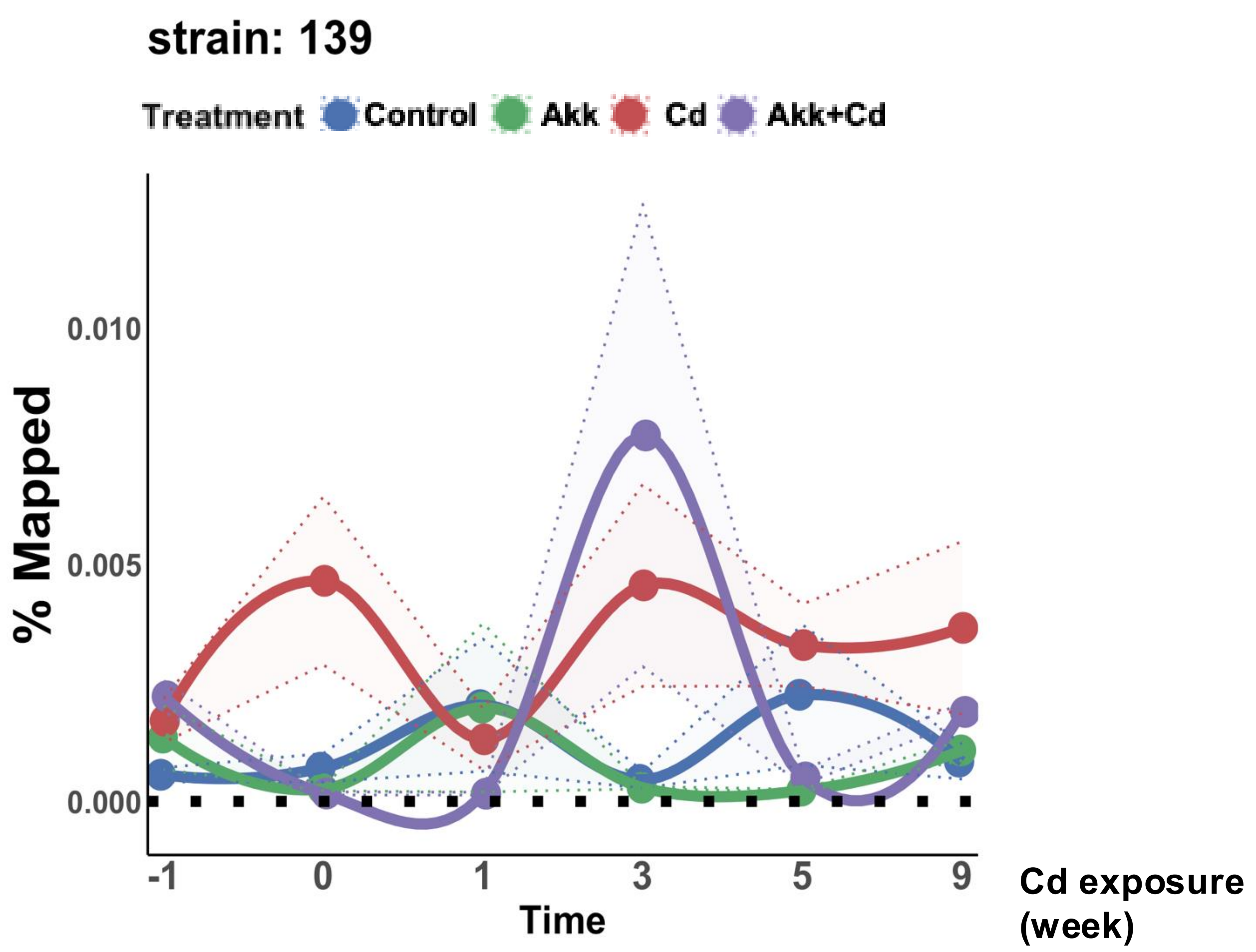


Supplementary Figure 1

Open field test (before treatment)

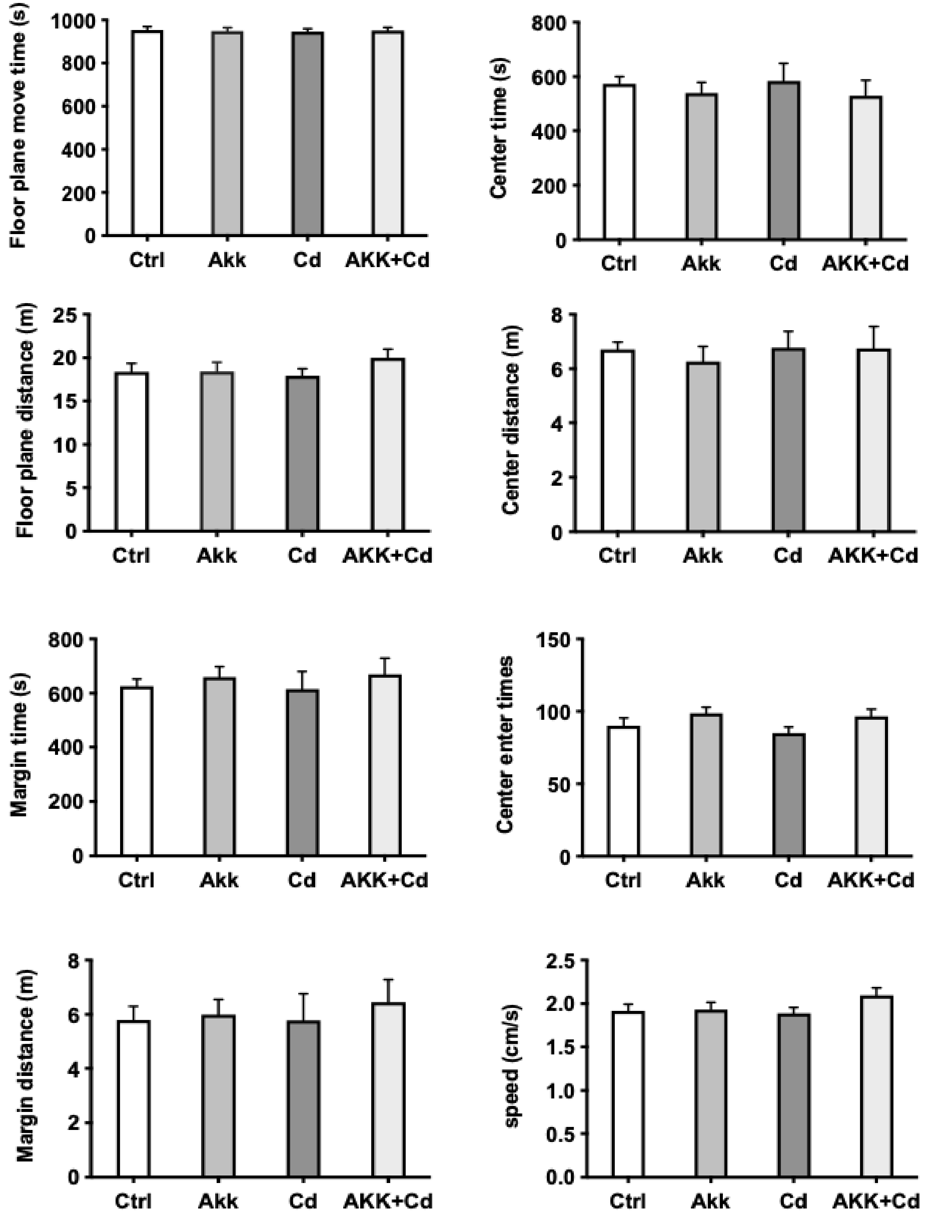


Open field test (after treatment)

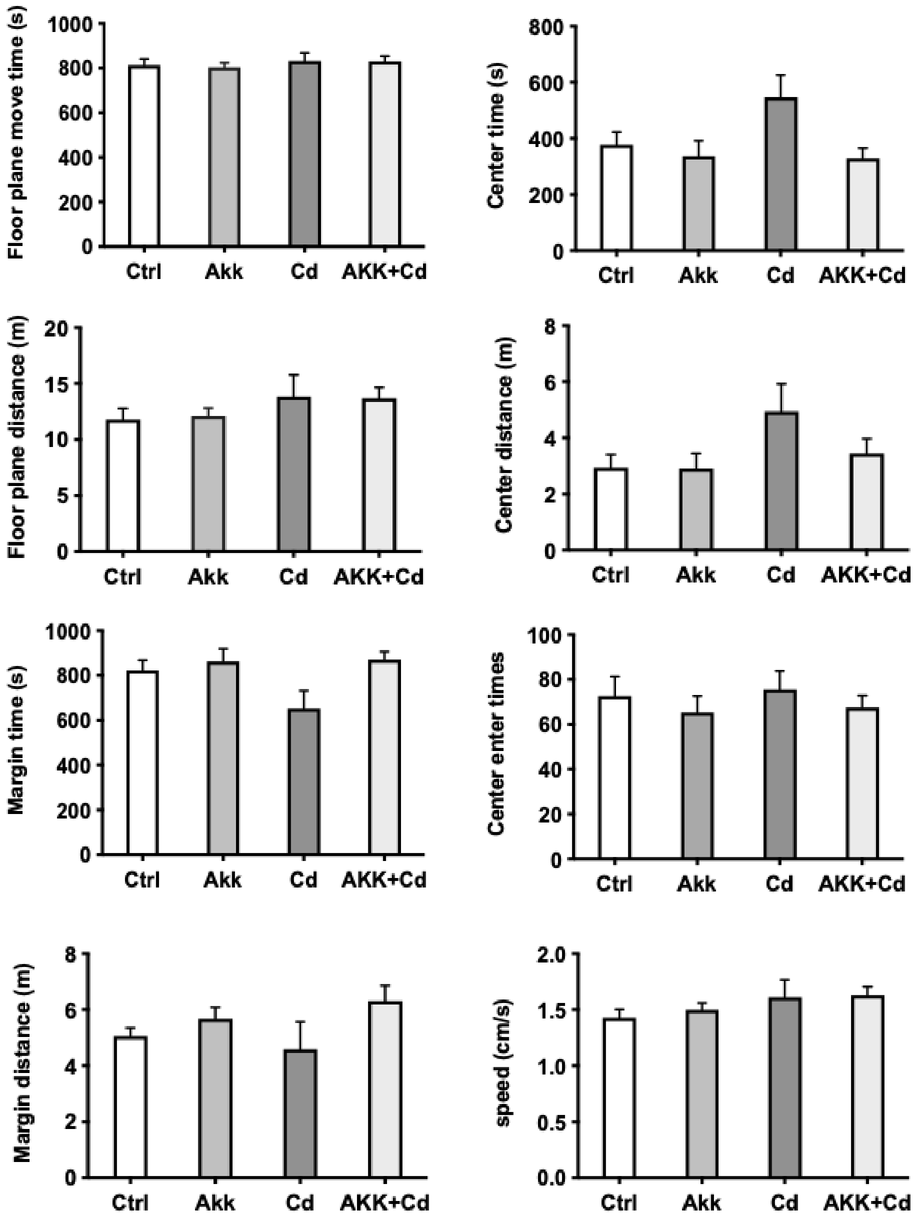


Supplementary Figure 2

### Supplementary Figure 3A

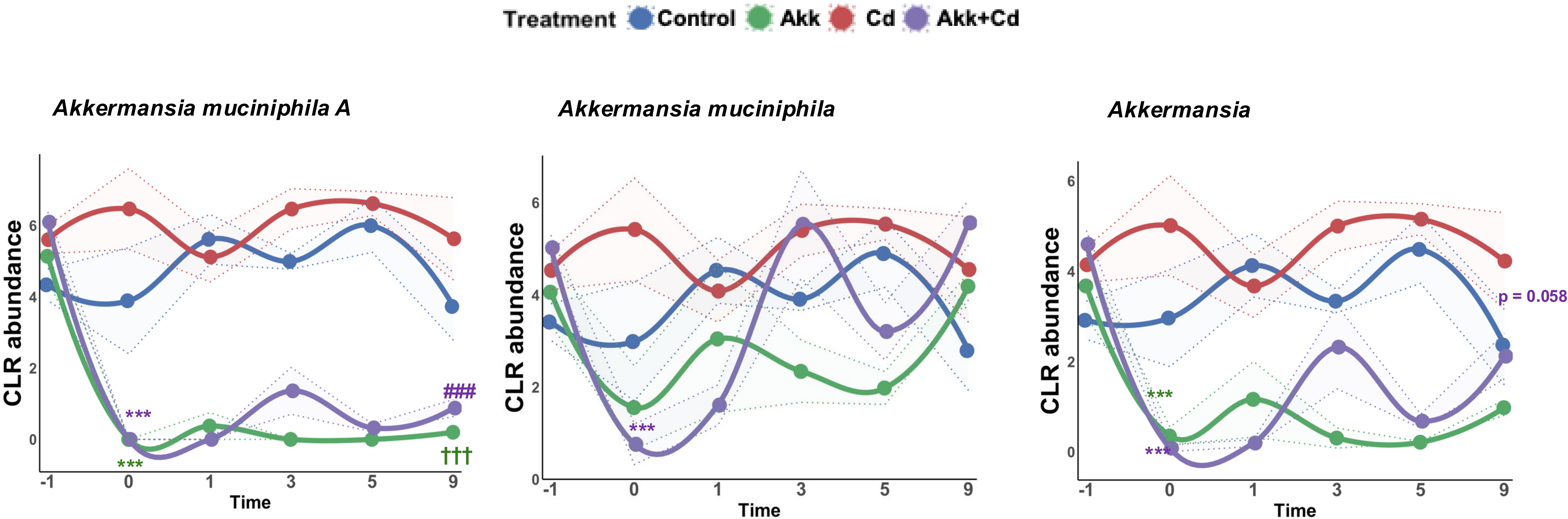

**Supplementary Figure 3B**

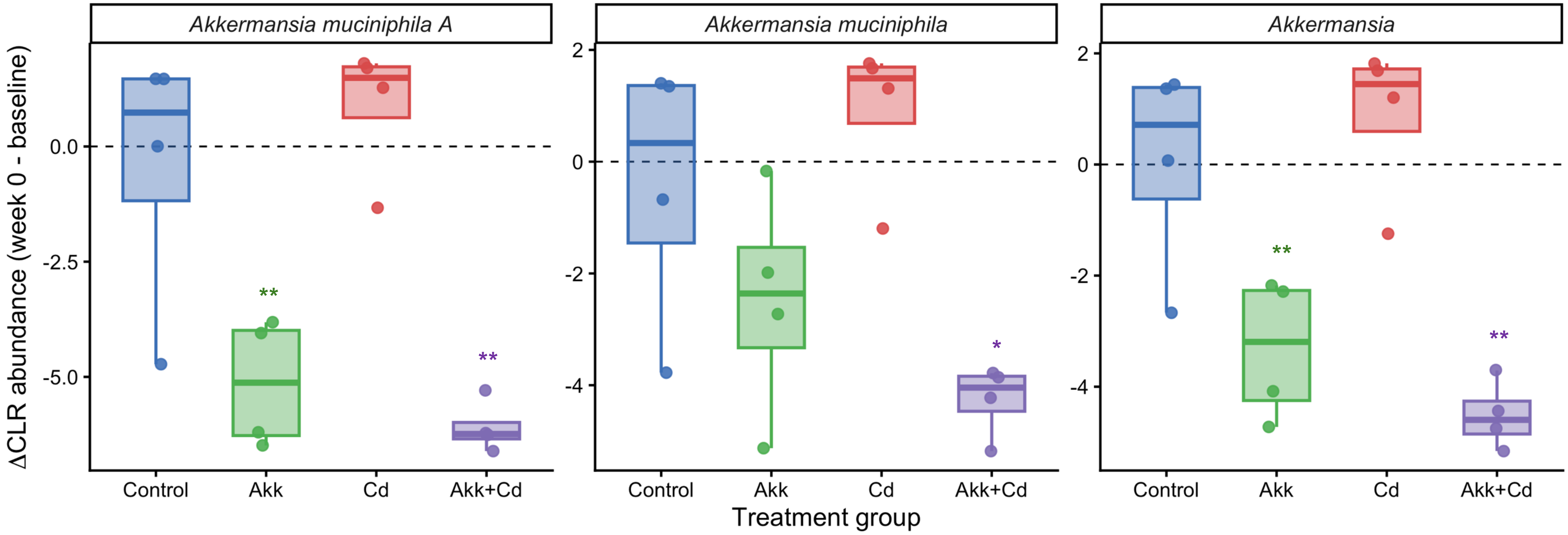

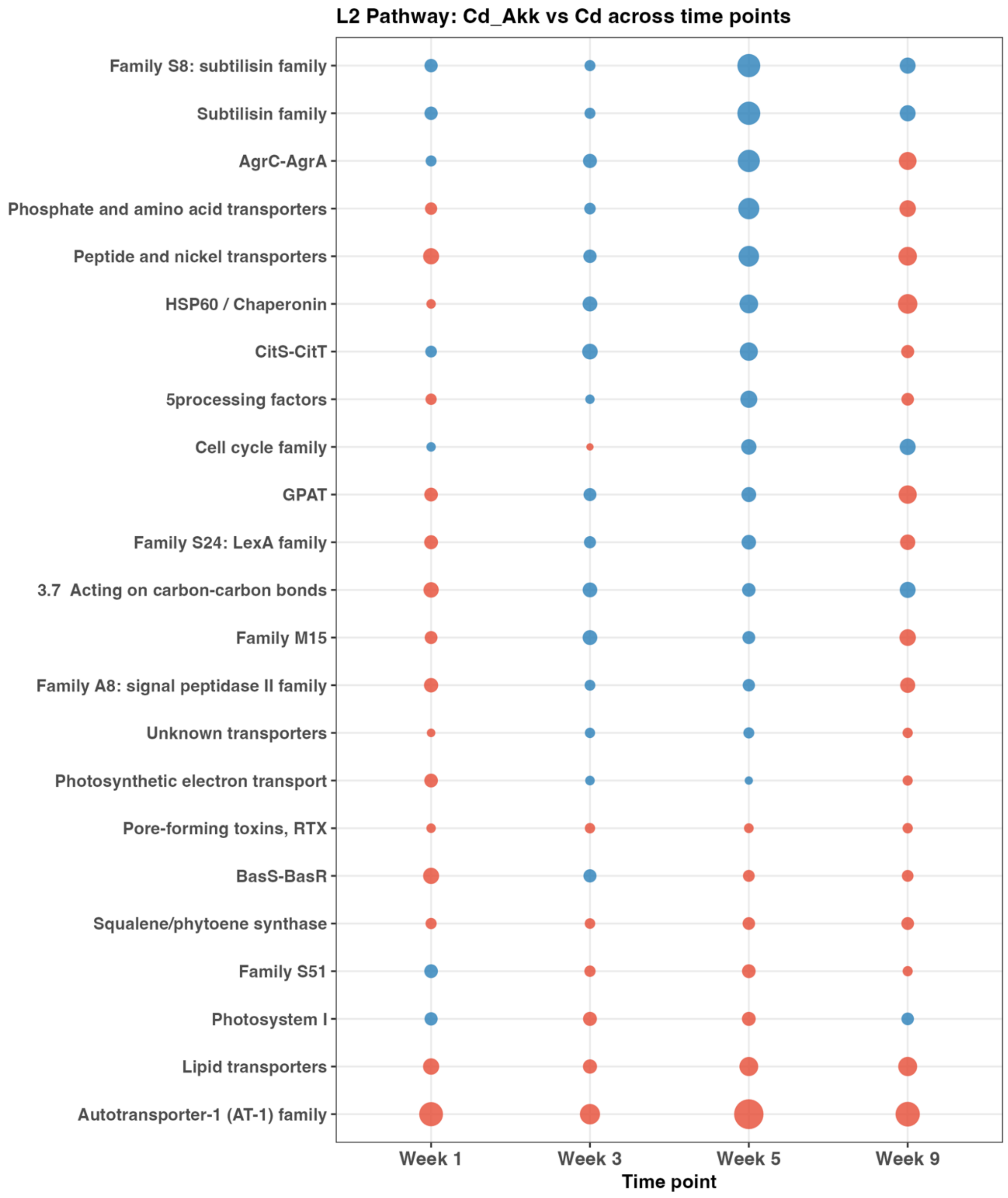


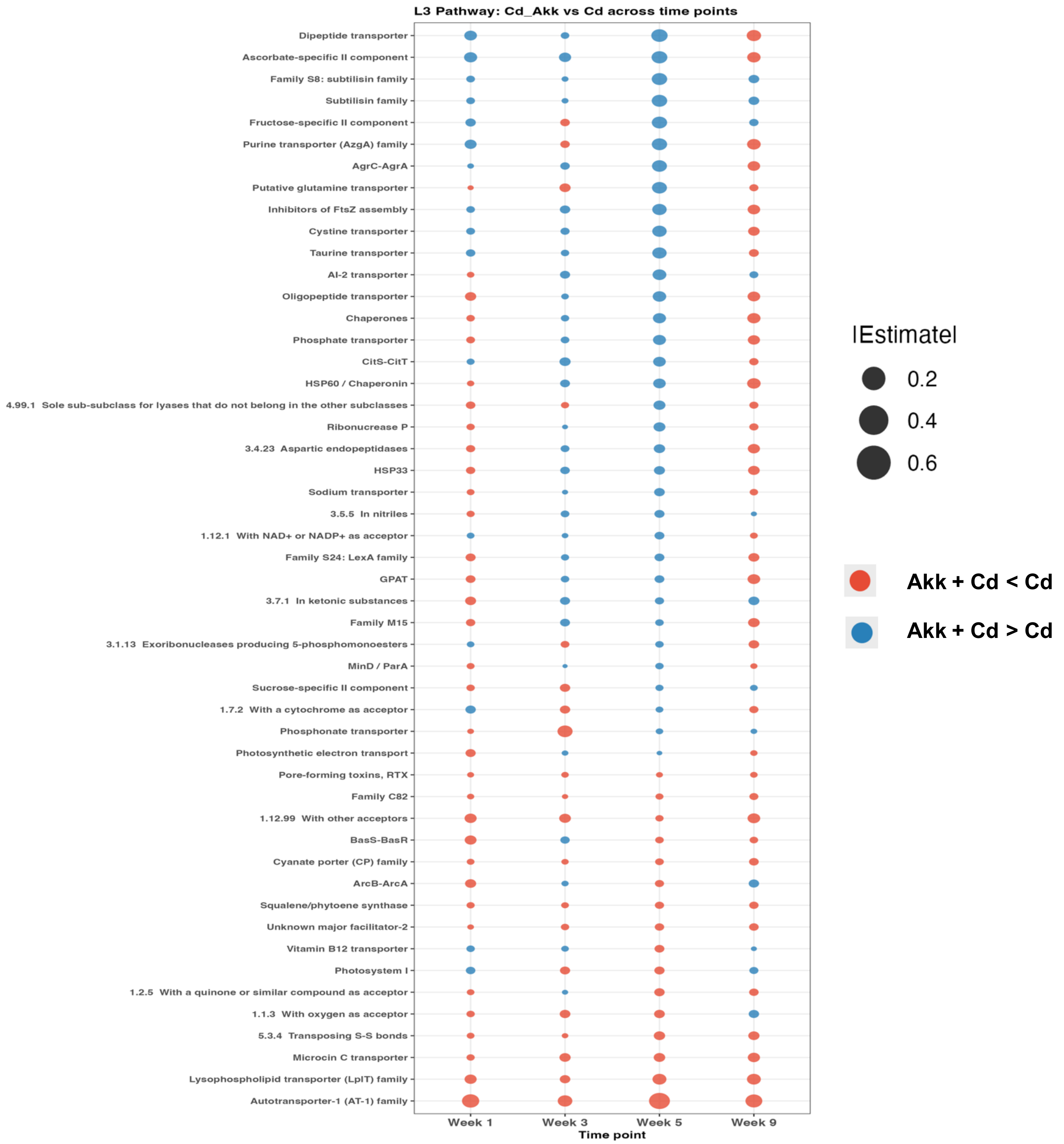


**Supplementary Figure 4**

**Supplementary Figure 5**

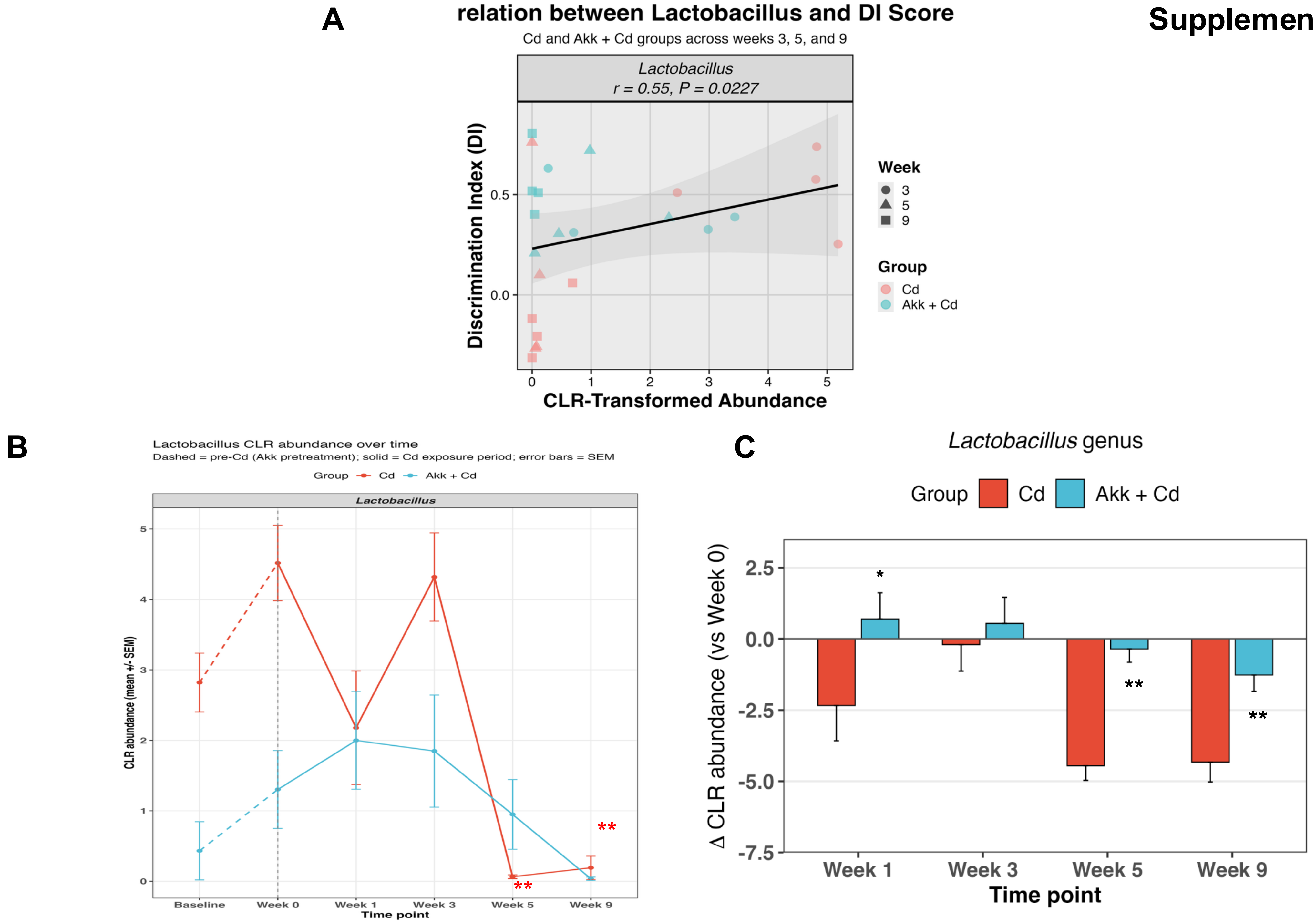

Serum

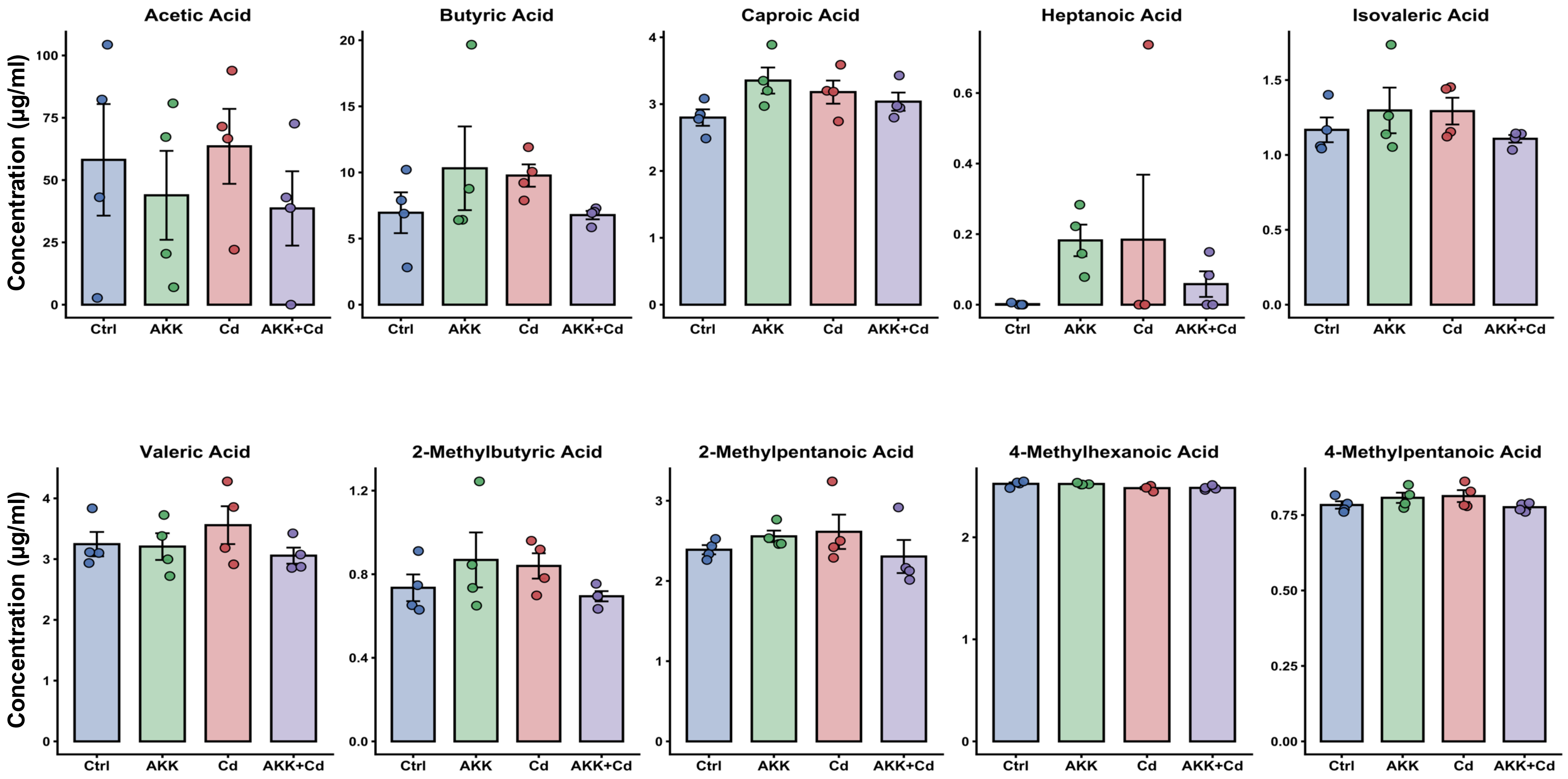


Supplementary Figure 6

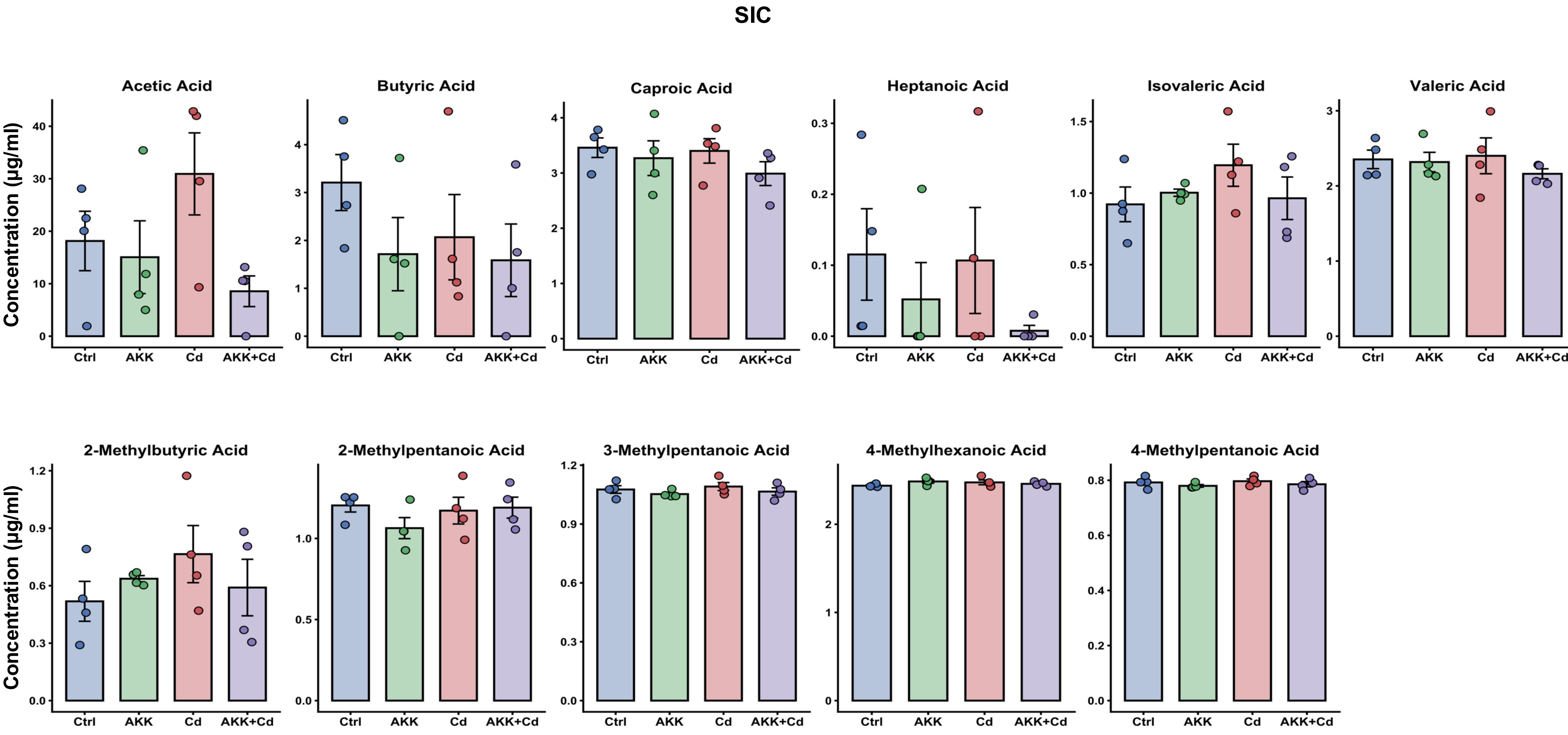


Supplementary Figure 7

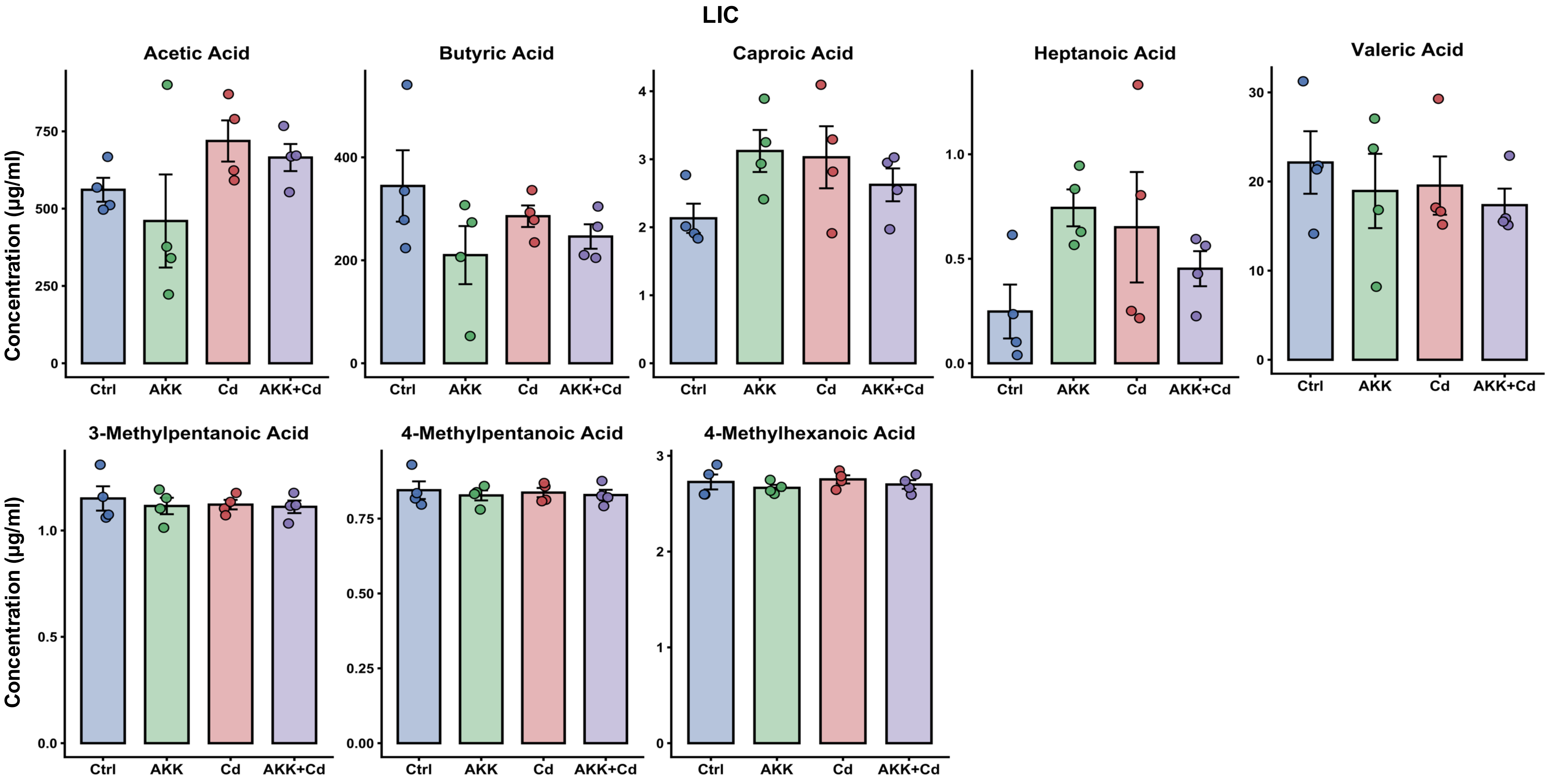


Supplementary Figure 8

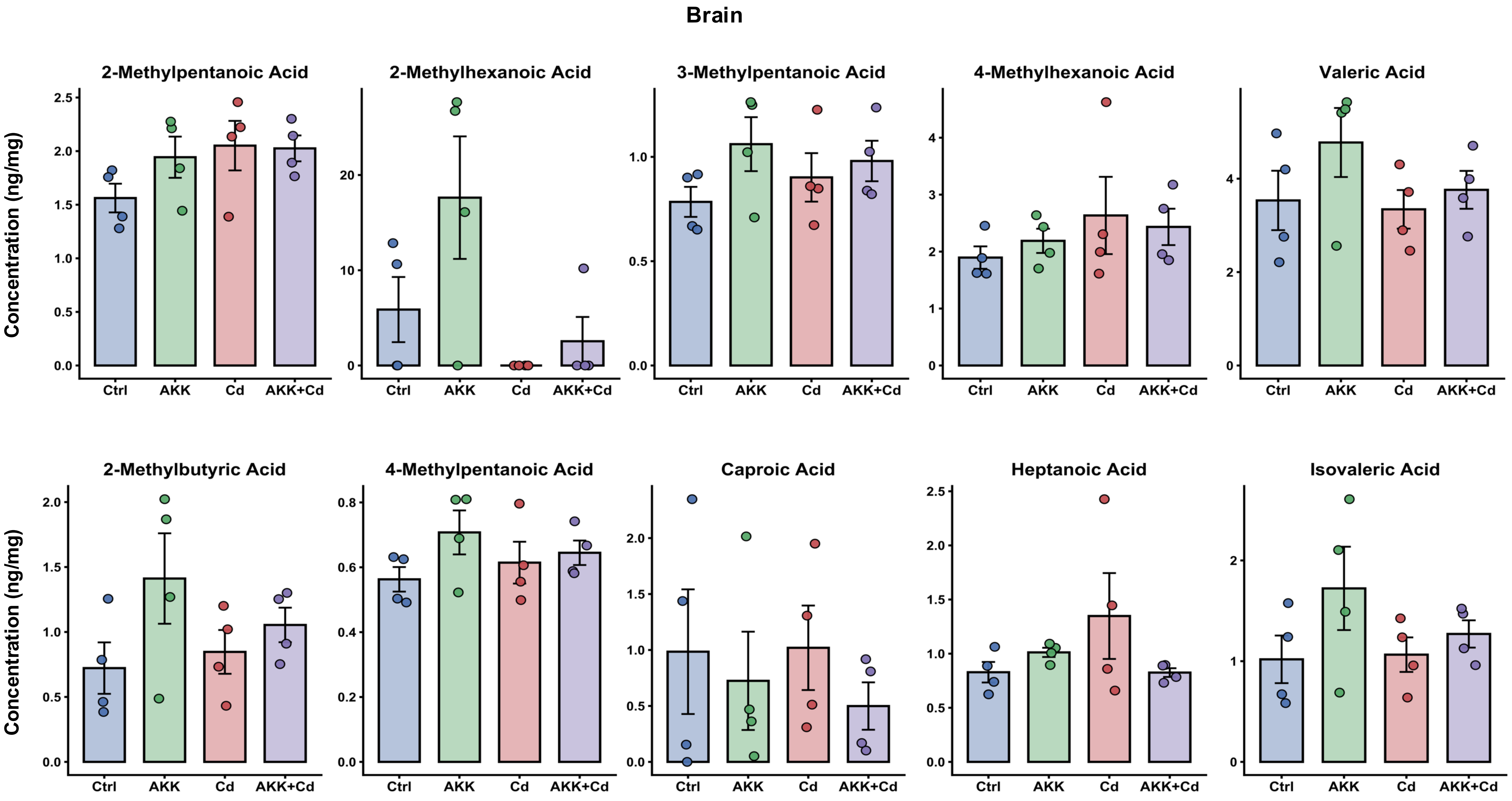


Supplementary Figure 9
